# SpRY greatly expands the genome editing scope in rice with highly flexible PAM recognition

**DOI:** 10.1101/2020.09.23.310839

**Authors:** Ziyan Xu, Yongjie Kuang, Bin Ren, Daqi Yan, Fang Yan, Carl Spetz, Wenxian Sun, Guirong Wang, Xueping Zhou, Huanbin Zhou

## Abstract

Plant genome engineering mediated by various CRISPR-based tools requires specific protospacer adjacent motifs (PAMs), such as the well-performed NGG, NG, NNG, *etc*., to initiate the target recognition, which notably restricts the editable range of the plant genome. In this study, we thoroughly investigated the nuclease activity and the PAM preference of two structurally-engineered SpCas9 variants (SpG and SpRY) in transgenic rice. Our study shows that SpG nuclease favors NGD PAMs, albeit less efficiently than the previously described SpCas9-NG and that SpRY nuclease achieves efficient editing across a wide range of genomic loci, exhibiting a preference of NGD as well as NAN PAMs. Furthermore, SpRY-fused cytidine deaminase hAID*Δ and adenosine deaminase TadA8e were generated, respectively. These constructs efficiently induced C-to-T and A-to-G conversions in the target genes toward various non-canonical PAMs, including non-G PAMs. Remarkably, high-frequency self-editing events (indels and DNA fragments deletion) in the integrated T-DNA fragments as a result of the nuclease activity of SpRY were observed, whereas the self-editing of SpRY nickase-mediated base editor was quite low in transgenic rice lines. In conclusion, the broad PAM compatibility of SpRY greatly expands the targeting scope of CRISPR-based tools in plant genome engineering.

## Background

CRISPR technologies, which enable a variety of precise genetic modifications through targeted genome editing across nearly all living organisms, have been intensively developed in recent years and revolutionized agricultural studies. CRISPR is derived from the defense system of bacteria and archaea, protecting them from the invasion of bacteriophages and mobile genetic elements [1]. A large collection of CRISPR/Cas systems have been identified by genome sequencing and metagenome studies. Among these, the type II CRISPR/SpCas9 from *Streptococcus pyogenes* has been extensively studied and adapted for genome editing. For example, SpCas9 nuclease can be used to genetically modify a specific target locus within the plant genome. Briefly, SpCas9 catalyzes a double-stranded break (DSB) which subsequently stimulates diverse DNA repair mechanisms [non-homologous end joining (NHEJ), microhomology-mediated end joining (MMEJ), and homology-directed repairs (HDR)] resulting in gene knockout, DNA fragment insertion, deletion, and replacement as specifically required [2-6]. Alternatively, the SpCas9(D10A) nickase can be utilized to drive various engineered nucleoside deaminases to the target region and catalyze the hydrolytic deamination of cytosines to uracils and/or adenosines to inosine within the editing window. As a result, the mismatched DNA base pairs are processed through the base excision repair pathway, resulting in various base transitions and transversions: C-to-D (where D is T, G, or A), G-to-H (where H is A, C, or T), A-to-G, and T-to-C [7-13]. Recently, the SpCas9(H840A) nickase-guided Moloney murine leukemia virus reverse transcriptase (M-MLV RT) has been developed in both rice and wheat, facilitating the introduction of all 12 base-to-base conversions, small insertion as well as deletions in a precise and targeted manner through reverse transcription of the pegRNA [14-17]. Thus, diverse customized CRISPR/SpCas9 tools for precise and versatile genome editing greatly accelerate our understanding of the genetic basis of economic traits and the generation of novel germplasms for crop breeding.

It’s well known that SpCas9 recognizes the canonical NGG PAM which is located immediately downstream of the target sequence [18]. Initial NGG PAM binding by SpCas9 triggers DNA strand separation locally and facilitates base pairing between spacer RNA and the target DNA strand [19]. In other words, the presence of a canonical NGG PAM near the target sites is critical for CRISPR/SpCas9-mediated genome editing. SpCas9 and its related tools have been shown to perform efficiently in genetic manipulation of a wide range of plant species [2, 10, 20-23]. However, the specific PAM requirement for SpCas9 recognition strongly restricts the targetable loci of the CRISPR/SpCas9-based editing tools, especially base editors because the targeted point mutation needs the availability of a PAM appropriately positioned [7-9, 24].

To expand the genome-targeting scope of CRISPR tools, many efforts have been focused on searching for new Cas variants and orthologs with novel PAM preferences. For example, directed evolution and structure-guided design of SpCas9 resulted in SpCas9-VRQR, xCas9, Cas9-NG variants that recognize non-canonical NGA and NG PAM sites in plant [25-28]. Multiple naturally occurring SpCas9 orthologues have been identified from *Streptococcus canis* (ScCas9), *Staphylococcus aureus* (SaCas9), *Streptococcus thermophiles* (St1Cas9), *Brevibacillus laterosporus* (BlatCas9) and have been demonstrated to edit plant genomic loci bearing NNG, NNGRRT, NNAGAAW, NNNCND PAM sequence, respectively [29-32]. Also, the type V Cas12a and Cas12b from diverse bacterial sources, which are distinct from Cas9, have been characterized with AT-rich PAM specificity and utilized successfully in targeted plant genome editing [33, 34]. Regardless of the fact that these Cas proteins with different PAM sequences significantly improve the targeting range, there are still many agronomic trait-related loci inaccessible for genome editing.

Very recently, two variants (SpG and SpRY) engineered from SpCas9-VRQR through structure-guided design have been reported to recognize a wider range of PAM sequences. SpG has higher genome-editing activity toward NG PAM than SpCas9-NG, whereas SpRY is capable of targeting almost all PAMs (NRN>NYN) [35]. Both SpG and SpRY exhibit robust activities with minimal side effects on a wide range of sites in human cells [35]. Furthermore, an evolved adenosine deaminase TadA8e, which catalyzes DNA deamination up to ∼1100-fold faster than the previous version TadA7.10, has been reported to substantially improve adenine base editing in human cells [36, 37]. However, whether SpG, SpRY, and TadA8e can be utilized to improve genome editing, especially base editing, in plants remains unknown. In this study, the efficiency of both SpG and SpRY nucleases toward various PAMs and its application in both cytosine and adenine base editing was investigated in detail using transgenic rice callus. Our study shows that SpG recognizes NG PAM sequences, but it’s outperformed by SpCas9-NG in rice. However, albeit with self-targeting activity on transfer T-DNA sequence, SpRY surpasses SpCas9-NG since it is capable of achieving efficient cleavage by its nuclease activity, cytosine base editing with hAID*Δ as well as adenine base editing with TadA8e at more relaxed PAM sites (NRN, where R is G or A) in rice. Therefore, SpRY and TadA8e facilitate the future design of genome editing tools in plants and broaden the possible applications in agriculture and plant biology.

## Results

### SpCas9-NG nuclease outperforms SpG on the minimal NG PAM in transgenic rice lines

We first compared the DNA-cleavage capability and the PAM specificity of SpG nuclease to that of SpCas9-NG (which is well known for NG PAM recognition) in targeted genome editing in transgenic rice. *SpCas9* carrying mutations D1135L/S1136W/G1218K/E1219Q/R1335Q/T1337R was rice codon-optimized (Table S1) and used to replace the SpCas9-NG gene in the binary vector pUbi:SpCas9-NG [28], resulting in pUbi:SpG in which *SpG* is under the control of the maize ubiquitin 1 promoter (Fig. 1a). The rice U6 promoter-driven sgRNA expression cassette can be shuttled into both binary vectors through Gateway recombination reaction as previously described [28]. The nuclease activity of SpG and SpCas9-NG was tested side-by-side with the same *sgRNA*s targeting the endogenous *OsPAL5, OsGSK4, OsCERK1, OsETR2*, and *OsRLCK185* genes toward four types of NGN PAMs through *Agrobacterium*-mediated rice transformation, respectively (Fig. S1; Fig. S2). Target regions in T0 independent transgenic callus lines were PCR amplified and directly subjected to Sanger sequencing.

**Fig. 1.**
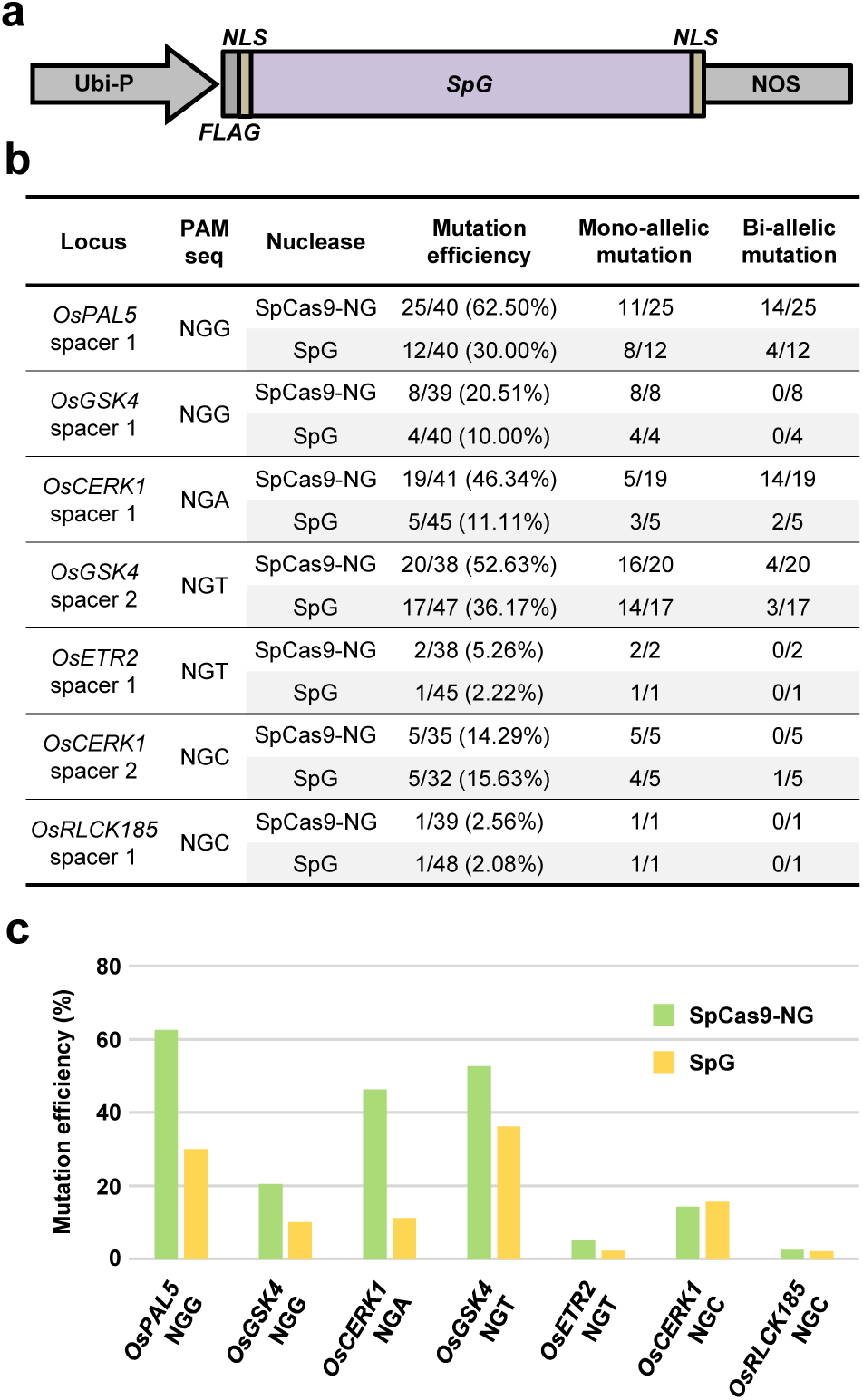
Analysis of SpG nuclease activity on different NGN PAMs in transgenic rice. **a** The gene construct of SpG nuclease used for genome editing in transgenic rice. Ubi-P, maize ubiquitin 1 promoter; *NLS*, nuclear localization sequence. **b** Summary of the genotyping results on T0 transgenic rice lines generated with SpG and SpCas9-NG nuclease. **c** Comparison of the mutation frequency by SpG and SpCas9-NG nucleases toward 4 NGN PAMs in T0 transgenic rice lines.

Analysis of 40 independent lines for each NGG PAM site tested with SpG nuclease revealed twelve lines that carried indel mutations in *OsPAL5* and four mutant lines of *OsGSK4*, representing 30% and 10% editing efficiency, respectively (Fig. 1b, c). By contrast, SpCas9-NG achieved 62.50% (25 out of 40 lines) editing efficiency in *OsPAL5*, a 1.1-fold increase over that of SpG; and 20.51% (8 out of 39 lines) editing efficiency in *OsGSK4*, a 1.1-fold increase over SpG (Fig. 1b, c). At the NGA PAM site tested, SpG generated much fewer indel mutations (< 4 fold) than SpCas9-NG, 11.11% of lines edited by SpG carried indels in *OsCERK1* as compared to 46.34% of the lines edited by SpCas9-NG (Fig. 1b, c). At the NGT PAM sites tested, the editing efficiency of SpG was 36.17% (17 out of 47 lines) in *OsGSK4* and 2.22% (1 out of 45 lines) in *OsETR2*; whereas the editing efficiency of SpCas9-NG was higher, being 52.63% and 5.26% in *OsGSK4* and in *OsETR2*, respectively (Fig. 1b, c). In the case of the NGC PAM, which has been reported to be less efficiently recognized by SpCas9-NG [28, 38], SpG edited *OsCERK1* and *OsRLCK185* at the corresponding sites with comparable efficiency to SpCas9-NG (Fig. 1b, c). It should be noted that both di-allelic and mono-allelic mutations were identified in T0 mutant lines generated by SpG toward all PAM sequences, being the ratios of mono-allelic edits relatively higher (Fig. 1b, c; Fig. S1; Fig. S2). Overall, these results indicate that the newly-engineered SpG nuclease is capable of recognizing minimal NG PAM in rice, albeit less efficiently than SpCas9-NG. Consequently, the activity of SpG in plants is different from the previously reported SpG-mediated genome editing efficiency in human cells [35].

### nuclease preferentially recognizes NAN and NGN PAM sequences in transgenic rice lines

Five other point mutations consisting of A61R, L1111R, N1317R, A1322R, and R1333P were further introduced into SpG to generate SpRY (Table S1; Fig S3a). Following the same experimental procedure mentioned above, the nuclease activity and PAM preference of SpRY were thoroughly investigated in transgenic rice calli. Thirty-two endogenous genomic sites bearing all 16 possible alternative PAM sequences, which vary at the second and the third positions, were used (Fig. S3-S16). Each transformation construct used harbored two *sgRNAs* targeting the same gene or different genes to the extent that multiplex genome editing and/or DNA fragment deletion could be achieved and investigated in rice cells.

Genotyping of all the T0 transgenic lines for the NGN PAM sites showed that SpRY induces indel mutations predominantly at the NGG PAM site (63.83% editing), followed by the NGA (> 74% editing) and NGT (> 47% editing) PAM sites. The NGC PAM site is the least recognized, achieving only a 2.13% editing efficiency (Fig. 2a, b). Thus, SpRY nuclease favors the recognition of NGD > NGC PAM in rice. On the NAN PAM sites tested, the editing efficiency of SpRY was highly variable, but it reached as high as 20.00% for a NAG PAM site in *OsMPK8*; 33.33% for a NAA site in *OsMPK9*; 51.28% for a NAT site in *OsMPK10*; and 29.79% for a NAC site in *OsCPK2* (Fig. 2a, b), suggesting that SpRY moderately recognizes NAN PAM sequences in rice, albeit in a locus-dependent manner. Across the rest 16 genomic sites with NYN PAMs, SpRY exhibited editing at 4 sites (25%), with high activity on a NTC PAM site in *OsCPK28* (73.91% efficiency) and weak activity on a NCG site in *OsCPK20* (2.94% editing), a NCT site in *OsMPK3* (4.26% editing), and a NCC site in *OsMPK4* (7.69% editing) (Fig. 2a, b). It should be mentioned that screening of all target genes revealed that the majority of edited lines contained mono-allelic mutations at the target sites, and that only 2 out of 47 independent lines (4.26% ratio) had genomic DNA fragment deletions in *OsCPK2* (Fig. 2a; Fig S3-S16). Combining all the data (Fig. 2b), we conclude that the structurally-engineered SpRY is capable of recognizing NRN and some NYN PAMs in rice, but with impaired nuclease activity as compared to that of SpCas9 reported previously [2].

**Fig. 2.**
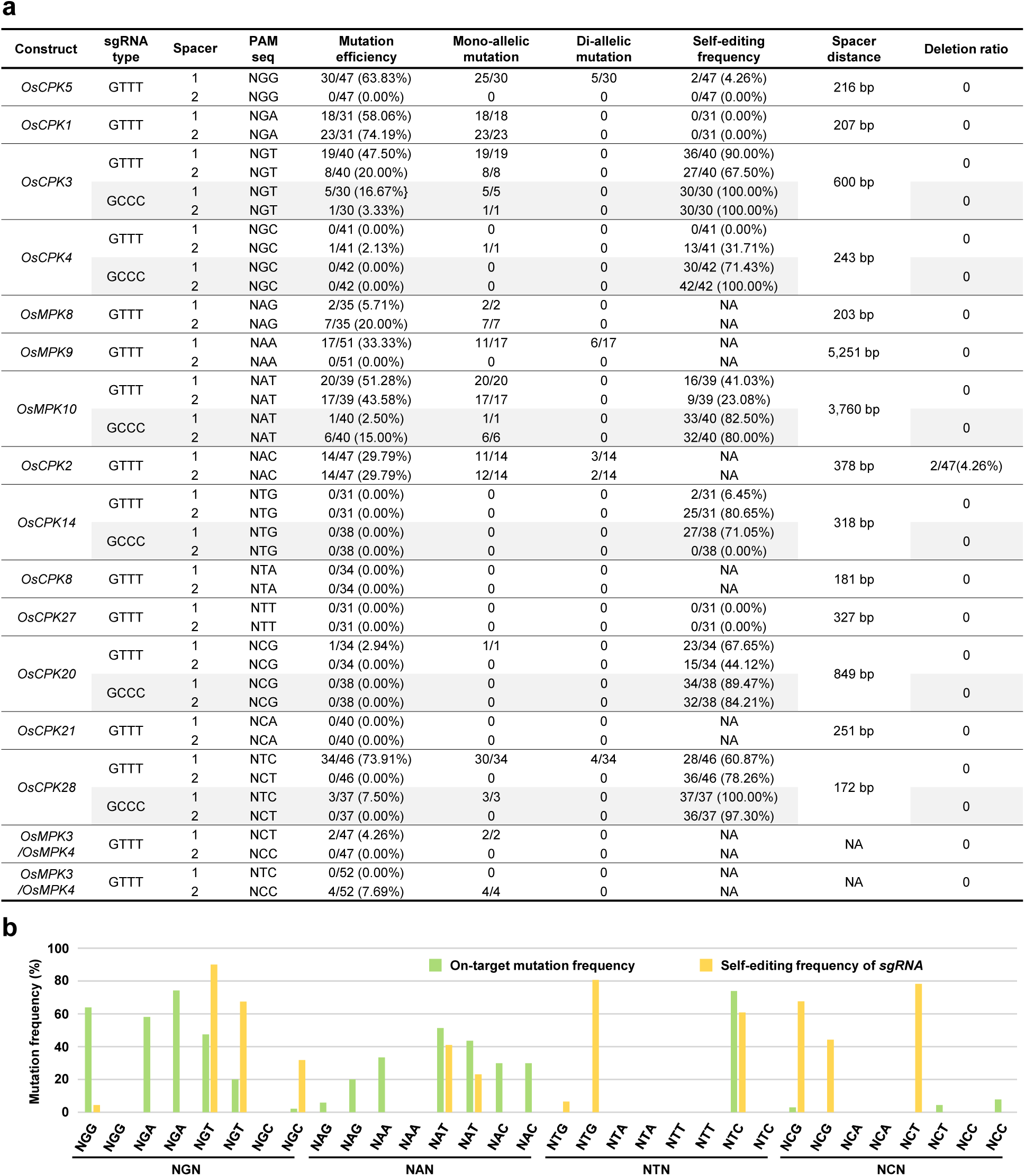
Analysis of SpRY nuclease activity on 16 possible NNN PAMs in transgenic rice. **a** Summary of the genotyping results on T0 transgenic rice lines generated with SpRY nuclease using different PAM sequences and sgRNA scaffolds. NA, not available. **b** Comparison of the on-target mutation efficiency and the self-editing frequency of *sgRNA* transgenes by SpRY nuclease toward different PAMs in T0 transgenic rice lines.

The broad PAM compatibility of SpRY nuclease implies that SpRY might be similar to SpCas9-NG which was reported to self-edit the single-guide RNA region of the transferred T-DNA in transgenic rice cells [39]. Therefore, the identities of the *sgRNA* transgenes in some *SpRY*-transgenic lines were examined by direct PCR sequencing. For the high-efficiency *OsCPK5-sgRNA1, OsCPK1-sgRNA1*, and *OsCPK1-sgRNA2*, no more than 4.26% lines were detected with modified *sgRNAs* (Fig. 2a, b). On the other hand, mutation frequencies at the other high-efficiency *sgRNA* sites (such as *OsCPK3-sgRNA1, OsCPK3-sgRNA2, OsMPK10-sgRNA1, OsCPK28-sgRNA1*) were much higher, ranging from 23.08% to 100% (Fig. 2a, b). Also, varying self-editing frequencies (0-80.65%) were observed in the low-efficiency *sgRNA*s (i.e. *OsCPK5-sgRNA2, OsCPK14-sgRNA1, OsCPK14-sgRNA2, OsCPK27-sgRNA1*) (Fig. 2a, b). It should be mentioned that on-target editing and self-editing of SpRY were detected simultaneously or alone in the individual transgenic line, and DNA fragment deletion events between two *sgRNAs* in the T-DNA regions were detected in targeting *OsCPK4* and *OsCPK28*, respectively (Fig. S6; Fig. S13). Combining all the data, we conclude that SpRY elicits complex self-editing events that might occur before and after gene editing of on-targets in transgenic rice.

A previous study has shown that utilizing a GCCC-type sgRNA scaffold alleviates the self-targeting property of SpCas9-NG without substantial loss of on-target activity in rice [39]. Therefore, we replaced the original GTTT-type sgRNA in our CRISPR/SpRY system with the GCCC-type sgRNA scaffold (Table S1). Subsequently, we tested the nuclease activity of our modified CRISPR/SpRY system in transgenic rice callus using the *sgRNAs* with high self-editing frequencies. To our surprise, replacing the GTTT-type scaffold with the GCCC-type scaffold did not show any improvement in on-target editing (Fig. 2a). On the contrary, the editing efficiency of *OsCPK3, OsMPK10*, and *OsCPK28* was significantly reduced (Fig. 2a). We further investigated the identities of each sgRNA transgene in all lines and found that, out of 12 sgRNAs examined, eleven lines carried mutations at high frequencies (Fig. 2a). Based on our data, we conclude that the GCCC-type sgRNA scaffold elicits stronger self-editing events in sgRNA transgenes, which results in a decrease in the efficiency of on-target editing events in rice, as compared to the naturally occurring (original) GTTT-type sgRNA scaffold.

### SpRY mediates cytosine base editing at non-canonical PAM sites in the rice genome

PAM preferences of Cas proteins largely limit the targeting scope of base editors [7-9, 28, 29]. Considering the greatly relaxed PAM compatibility of SpRY, we tested whether it could improve cytosine base editing in transgenic rice when fusing to the hyperactive cytidine deaminase hAID*Δ. The SpRY nuclease was mutated into the SpRY(D10A) nickase and employed to replace the *SpCas9n* gene in the previously-reported cytosine base editor rBE9 which enables C-to-D editing [7], resulting in rBE66 (Fig. 3a).

**Fig. 3.**
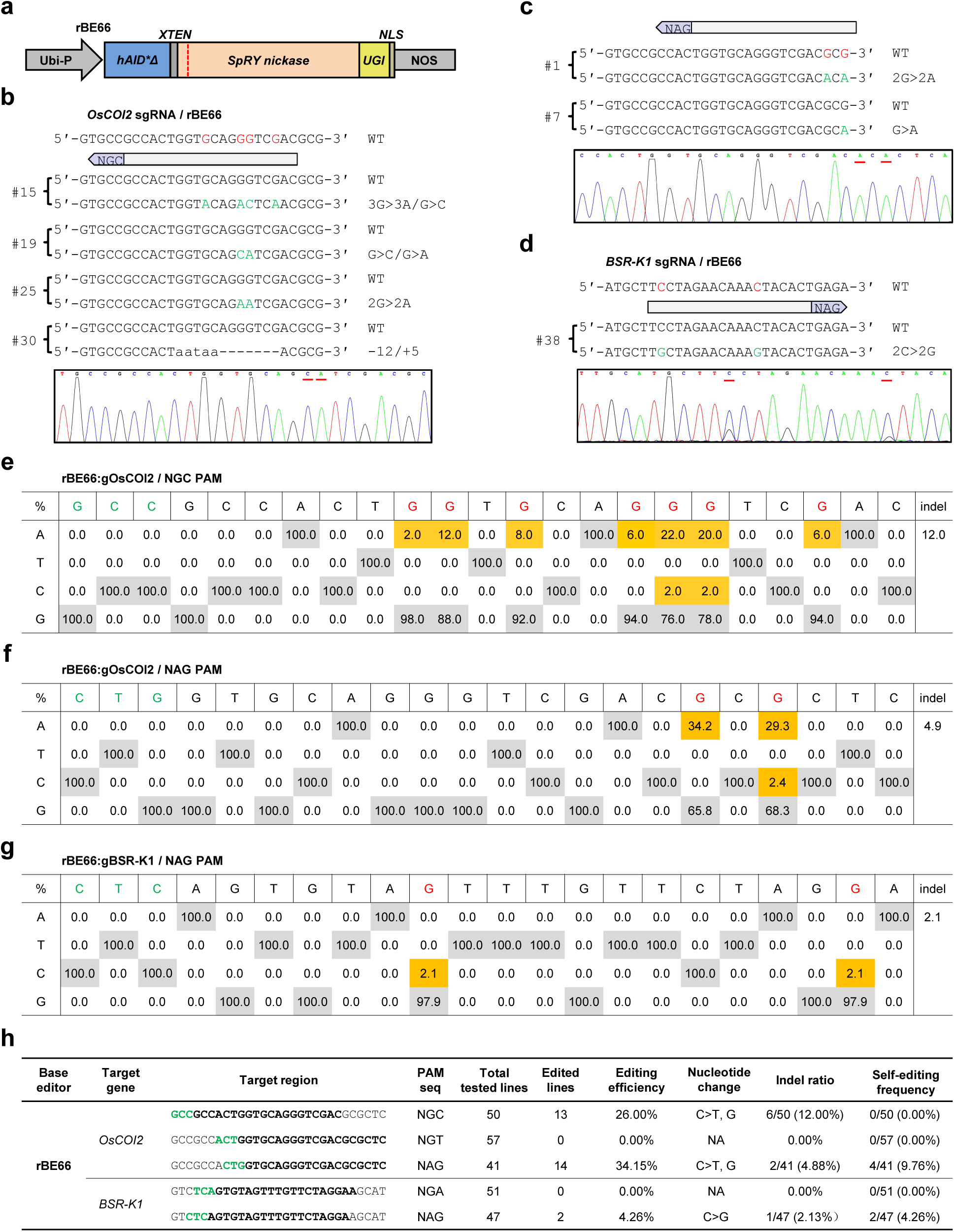
Efficient cytosine base editing mediated by SpRY-fused hAID*Δ toward non-canonical PAMs in rice. **a** The gene construct of *rBE66* used for cytosine base editing in transgenic rice. Ubi-P, maize ubiquitin 1 promoter; *hAID*Δ*, truncated version of hyperactive human *AID* gene; *UGI*, uracil glycosylase inhibitor; *NLS*, nuclear localization sequence. **b, c**, and **d** Representative edited alleles of *OsCOI2* toward a NGC PAM (**b**) and a NAG PAM (**c**), and *BSR-K1* toward a NAG PAM (**d**) generated by rBE66 in T0 transgenic rice lines. The PAM sequences for each target site are shown in the pale purple arrows; the candidate bases in the putative editing window for editing and detected nucleotide changes are highlighted in red and green, respectively; the nucleotide substitutions are underlined in the sequencing chromatograms. **e, f** and **g** Summary of nucleotide changes in the editing window of *OsCOI2* toward a NGC PAM (**e**) and a NAG PAM (**f**), and *BSR-K1* toward a NAG PAM (**g**) caused by rBE66 in T0 transgenic lines. The PAM sequences and the detected nucleotide changes are highlighted in green and red, respectively. **h** Summary of the genotyping results on T0 transgenic rice lines generated with rBE66. NA, not available.

The base-editing activity of rBE66 was then investigated in transgenic rice callus with three sgRNAs targeting the endogenous *OsCOI2* gene toward non-canonical NGC, NGT, and NAG PAMs, respectively. Intriguingly, of the 50 independent transgenic lines confirmed by sequencing, thirteen lines were identified with nucleotide mutations at the NGC PAM site which is presumably less efficient for SpRY targeting (Fig 3b, e, h). Among these thirteen lines, twelve lines (24% efficiency) carried nucleotide substitutions (predominantly the C-to-T conversion) in the typical editing window whereas six lines (12% frequency) carried indels (Fig 3e). These indels likely result from both the deaminase activity of hAID*Δ and the nickase activity of SpRY based on the position of the indels. For the other two PAMs tested in *OsCOI2*, while the NGT PAM was inefficient, the 1 bp-shifted NAG PAM resulted in a base editing efficiency of 34.15% (14 out of 41 lines) and indel frequency of 4.88% (2 out of 41 lines) (Fig 3c, f, h). rBE66 was also tested with two sgRNAs targeting the endogenous *BSR-K1* gene at NGA and NAG PAM sites, respectively. After genotyping the transgenic lines obtained for both transformation constructs, we only identified a single heterozygous line (2.13%) with base editing events and a single heterozygous line (2.13%) carrying indel at the NAG PAM site (Fig 3d, g, h). Finally, the self-editing effect of rBE66 was also investigated. We identified 4 lines (9.76% ratio) and 2 lines (4.26% ratio) carrying nucleotide substitutions in *OsCOI2-sgRNA* (NAG PAM) and *BSR-K1-sgRNA* (NAG PAM) transgenes, respectively (Fig. 3h, Fig. S17a). These data, combined, indicate that SpRY is compatible with hAID*Δ, greatly expanding the targeting scope of hAID*Δ-mediated cytosine base editor in rice. Thus, SpRY has very promising prospects in targeted base editing in plants due to the broad PAM compatibility and negligible self-editing effect.

### SpRY-fused TadA8e monomer enables efficient adenine base editing at non-G PAM sites in the rice genome

The CRISPR/Cas-guided adenine deaminase heterodimer TadA:TadA7.10 is capable of introducing A-to-G conversion in rice [9, 28, 29]. Very recently, an evolved version TadA8e with a more robust enzymatic activity has been characterized in human cells [36, 37]. Therefore, we investigated whether SpRY could improve adenine base editing in rice by fusing it to TadA8e. *TadA7*.*10* carrying mutations A109S/T111R/D119N/H122N/Y147D/F149Y/T166I/D167N was rice codon-optimized (Table S1) and fused to the 5’-terminus of *SpRY(D10A)*. The chimeric gene was utilized to replace the previously-reported adenine base editor gene *rBE14* in the binary vector pUbi:rBE14 [9], resulting in pUbi:rBE62 (Fig. 4a).

**Fig. 4.**
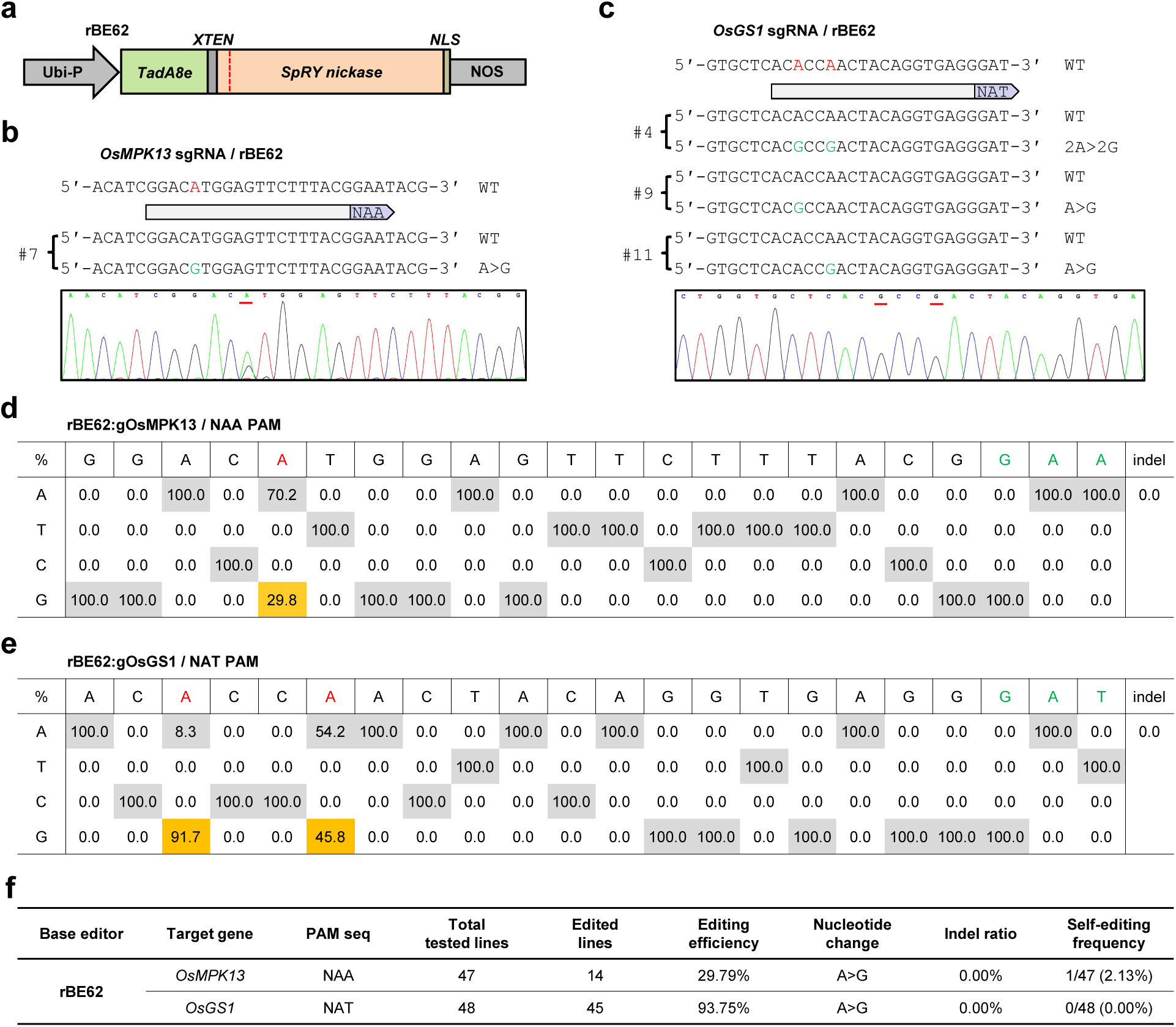
Efficient adenine base editing mediated by SpRY-fused TadA8e monomer toward non-G PAMs in rice. **a** The gene construct of *rBE62* used for adenine base editing in transgenic rice. Ubi-P, maize ubiquitin 1 promoter; *TadA8e*, the evolved *E*.*coli TadA7*.*10* gene; *XTEN*, 16 amino acid flexible linker; *NLS*, nuclear localization sequence. **b** and **c** Representative edited alleles of *OsMPK13* (**b**) and *OsGS1* (**c**) generated by rBE62 in T0 transgenic rice lines. The PAM sequences for each target site are shown in the pale purple arrows; the candidate bases in the putative editing window for editing and detected nucleotide changes are highlighted in red and green, respectively; the nucleotide substitutions are underlined in the sequencing chromatograms. **d** and **e** Summary of nucleotide changes in the editing window of *OsMPK13* (**d**) and *OsGS1* (**e**) caused by rBE62 in T0 transgenic rice lines. The PAM sequences and the detected nucleotide changes are highlighted in green and red, respectively. **f** Summary of the genotyping results on T0 transgenic rice lines generated with rBE62. The PAM sequences and the target regions are highlighted in green and bold, respectively.

The endogenous *OsMPK13* and *OsGS1* gene were utilized to test rBE62’s activity toward the NAA and NAT PAM in transgenic rice calli, respectively. Strikingly, 14 out of 47 lines for the NAA PAM and 45 out of 48 lines for the NAT PAM were identified with a pure A-to-G conversion at the target sites, representing 29.79% and 93.75% adenine editing efficiency, respectively (Fig 4b-f). All mutant lines carried mono-allelic mutations and no indel mutations were detected (Fig 4b, c). Furthermore, we genotyped the *sgRNA* transgenes in all lines to evaluate the self-targeting effect of rBE62. Only one self-edited line (2.13% ratio) in which the *sgRNA* transgene carried A-to-G conversion was detected (Fig S17b). Nevertheless, no modification was observed in the target region since the entire *OsMPK13* was intact. These data, combined, indicate that the adenine base editor rBE62 (engineered with SpRY and TadA8e monomer as mentioned above) is capable of efficiently inducing A-to-G conversions in a wide editable range of the rice genome, including non-G PAM sites. To our knowledge, this is the first time a base editor which recognized non-G PAM sites in plants is described.

## Discussion

PAM preference is the key limitation to each CRISPR-based tool for targeted genome editing, especially for base editing since it requires the precise positioning of Cas proteins at a given site. Therefore, reducing or eliminating the PAM requirement of Cas proteins will substantially advance various CRISPR technologies. To date, several other Cas proteins with altered or relaxed PAM specificities besides SpCas9, including SpCas9-VQR (NGA), SpCas9-VRER (NGA), SpCas9-NG (NG), ScCas9 (NNG), and SaKKH-Cas9 (NNNRRT), have been successfully adopted in both cytosine and/or adenine base editing in plants [8, 28, 29, 38]. The latest reported SpCas9 variants, SpG and SpRY with relaxed PAM specificity [35], might further optimize plant CRISPR tools for targeted base editing. However, different from the original report that SpG nuclease shows high-activity editing across all NGN PAM sites in human cells [35], we observed a different PAM preference (NGN/NAN/NTN>NCN) in transgenic rice lines. Moreover, our data also shows that the editing efficiency of SpG is lower than that of SpCas9-NG in transgenic rice lines. Therefore, we believe that SpCas9-NG is still the best genome editing player for NGN PAM sites in plants.

SpRY exhibits a similar preference for NGN PAM sequences as SpG. However, SpRY nuclease is also capable of processing many NAN, NTC, NCT, NCC, NCG PAM sites with varying efficiency, which have previously been inaccessible with CRISPR tools. SpRY thereby greatly increases the number of targeting sites previously not available for genome editing in rice. However, the broadened PAM compatibility of SpRY nuclease can also result in complex and unpredictable self-targeting of sgRNA transgenes as shown in our study, which presumably affects the on-target editing at both early- and late-transformation-stage. Nevertheless, we show that the naturally occurring GTTT-type of guide RNA scaffold outperforms the modified GCCC-type [39] when using SpRY to edit some endogenous genomic sites, and we do not recommend SpRY for multiple gene knocking out since it causes loss of sgRNA transgenes in some cases. More GTN- and GCN-type sgRNA scaffolds in self-editing and the heritability of edited genes in generations should be investigated in the future.

The SpRY nickase was compatible with both hAID*Δ and TadA8e deaminases and achieved efficient cytosine and adenine base editing at various non-canonical PAM sites, especially non-G PAM sites. Intriguingly, compared to the high-frequency self-editing caused by the SpRY nuclease in transgenic rice, significantly low ratios of self-editing of *sgRNA* transgenes was detected in our base editing assay, in which mutations are the result of the activity of the TadA8e deaminase and not the SpRY nickase. In our study, we also show that the newly evolved TadA8e, as reported in human cells [36], enables efficient A-to-G conversion in rice as well. Therefore, both SpRY and Tad8e can be presumably utilized to multiplex base editing in rice and further optimize other plant base editors in terms of targeting scope and editing efficiency.

The broadened PAM compatibility of SpRY raises the possibility of increased off-target editing across all plant genomes. To address this question, the fidelity-enhancing substitutions of SpCas9-HF1 could be introduced into the SpRY nuclease to alleviate its off-target effect in rice as reported in human cells [35]. Alternatively, careful design of the sgRNA is another way to avoid off-target cleavage. For example, off-target mutations were detected only for *sgRNAs* targeting *OsCPK3* and *OsCPK28* in our study (Fig. S18). Regarding the SpRY nickase-mediated base editing with specific sgRNAs, hAID*Δ and TadA8e-dependent off-target base editing are the major issues, which might be minimized through protein engineering in the future. Nevertheless, timely isolation of the transgene-free, gene-edited lines in T1 progenies to prevent continual editing is highly recommended, and backcrossing it with the recipient material to purify the genetic background can be carried out if needed.

## Conclusions

Overall, this article is the first report describing efficient base editing at minimal NA PAM sites in the plant genome. The structurally-engineered SpRY efficiently recognizes NAN in addition to NGN PAM sequences, greatly expanding the targetable range of the rice genome. With the new SpRY-based tools added into the toolbox, plant biologists can easily carry out high-resolution genome editing now, and we recommend using SpCas9 for target sites carrying NGG PAMs, SpCas9-NG for NGH PAMs (where H is A, C, or T), ScCas9 for NHG PAMs, and SpRY for NAH PAMs.

## Methods

### Rice materials

Rice cultivars Kitaake (*Oryza sativa L*. ssp. *Geng*) was used in this study and kept in our lab. Rice plants were cultivated in the paddy field under natural conditions in normal rice growing seasons and immature seeds were harvested from expanded panicles for use in rice transformation.

### Vector design and plasmid construction

The coding region of *SpG* gene [35] flanked by two nuclear localization signal (*NLS*) sequences was rice codon-optimized, the 5’ region (*SpG-fg1*), and the 3’ region (*SpG-fg2*) were separately synthesized by Tsingke (Beijing) (Table S1). The full-length *SpG* was assembled using the overlapping-extension PCR-based method with the high-fidelity DNA polymerase Phusion (NEB) and the primers listed in Table S3, and directly cloned downstream of the maize ubiquitin 1 promoter and upstream of the *NOS* terminator by replacing the *Cas9-NG* gene in pUbi:Cas9-NG [28] through BamHI/SpeI digestion and DNA ligation, resulting in the binary vector pUbi:SpG.

For the construction of the binary vector of SpRY nuclease, a point mutation of GCG to AGG (A61R) was first introduced into *SpG-fg1* by PCR amplification of the whole plasmid pUC57:SpG-fg1 with the primer pairs SpRY-F1/SpRY-R1. The amplicon was self-ligated and confirmed by sequencing, resulting in pUC57:SpRY-fg1. Next, the 3’ region of *SpRY* [35] fused with an *NLS* sequence was rice codon-optimized (*SpRY-fg2*) and subjected to synthesizing by Tsingke (Beijing) (Table S1). Finally, both *SpRY-fg1* and *SpRY-fg2* were assembled and pUbi:SpRY was constructed in the same manner as mentioned for *SpG*.

For the construction of cytosine adenine base editors, the gene fusion of *hAID*Δ, SpRY*, and *UGI* fragment was carried out with a simple PCR-based cloning strategy. Briefly, the full-length *SpRY* gene was PCR amplified using the primer pairs OsCas9-Fg1-F1/OsCas9-Fg1-R1 and pUbi:SpRY as the template, and ligated with the backbone of pUC19:rBE9 [7] carrying both *hAID*Δ* and *UGI* fragment, which was amplified with the primer pairs UGI-F1/rAPO-R1, resulting in pUC19:rBE66. For the construction of adenine base editors, the *TadA8e* fragment was codon-optimized and subjected to synthesizing by Tsingke (Beijing) (Table S1), and fused to the 5’ end of *SpRY* using the overlapping-extension PCR-based method with the primers listed in Table S3, resulting in *rBE62* fragment. Finally, rBE66 and rBE62 were cloned into the binary vector by BamHI/SpeI digestion as described above, resulting in pUbi:rBE66 for cytosine base editing and pUbi:rBE62 for adenine base editing in rice, respectively.

For the construction of guide RNA gene, the GCCC-type sgRNA [39] was subjected to synthesizing by Tsingke (Beijing) (Table S1), amplified with the primer pairs gRNA4-NG-F2/gRNA4-NG-R2, and used to replace the GTTT-type sgRNA in pENTR4:sgRNA4 using In-Fusion HD Enzyme Premix (Clontech), resulting in the entry vector pENTR4:sgRNA4-NG.

For the construction of the final T-DNA transformation plasmids, the 20-bp complementary oligos (Table S2), corresponding to each target site (Table S3) and carrying appropriate 4-bp adaptor, were phosphorylated, annealed, and inserted into the BsaI-digested pENTR4:sgRNA4 or pENTR4:sgRNA4-NG. Each sgRNA expression cassette was then shuffled into the respective binary vector by LR clonase (Invitrogen). All primers used here are listed in Table S3.

The identities of *SpG, SpRY, rBE66*, and *rBE62* in vectors and oligo insertions in each construct were confirmed by Sanger sequencing.

### *Agrobacterium*-mediated rice transformation

T-DNA transformation plasmids harboring gene-targeting *sgRNA* were transferred into the *A. tumefaciens* strain EHA105 by electroporation. Rice callus derived from immature seeds of Kitaake were used for stable transformation following a protocol as described previously [24].

### Genotyping of transgenic rice lines

Genomic DNA was isolated from each T0 transgenic callus line using the hexadecyltrimethylammonium bromide (CTAB) method [40]. PCR amplification of the targeted genomic region was carried out with specific primers listed in Table S3, and the PCR products were subjected to Sanger sequencing. Besides, a selection of PCR products with low quality of sequencing data was cloned into the pEASY-Blunt cloning vector (TransGen Biotech), and colonies with insertion were randomly chosen for Sanger sequencing.

## Acknowledgements

We thank Sujie Zhang and Xueqi Li for assistance with tissue culture of rice and DNA manipulation.

## Authors’ contributions

HZ, XZ, GW, CS, SW, and BR designed the research; ZX, YK, BR, DY, and FY conducted the experiments and analyzed the data; HZ, XZ, and GW supervised the research; HZ, XZ, and CS wrote the original draft; all authors participated in discussion and revision of the manuscript.

## Ethics approval and consent to participate

Not applicable.

## References

1. Hille F, Richter H, Wong SP, Bratovic M, Ressel S, Charpentier E: The biology of CRISPR-Cas: backward and forward. Cell 2018, 172:1239–1259.

2. Zhou H, Liu B, Weeks DP, Spalding MH, Yang B: Large chromosomal deletions and heritable small genetic changes induced by CRISPR/Cas9 in rice. Nucleic Acids Res 2014, 42:10903–10914.

3. Li J, Meng X, Zong Y, Chen K, Zhang H, Liu J, Li J, Gao C: Gene replacements and insertions in rice by intron targeting using CRISPR-Cas9. Nat Plants 2016, 2:16139.

4. Lu Y, Tian Y, Shen R, Yao Q, Wang M, Chen M, Dong J, Zhang T, Li F, Lei M, Zhu J-K: Targeted, efficient sequence insertion and replacement in rice. Nat Biotechnol 2020, DOI: 10.1038/s41587-020-0581-5.

5. Tan J, Zhao Y, Wang B, Hao Y, Wang Y, Li Y, Luo W, Zong W, Li G, Chen S, et al: Efficient CRISPR/Cas9-based plant genomic fragment deletions by microhomology-mediated end joining. Plant Biotechnol J 2020, DOI: 10.1111/pbi.13390.

6. Sun Y, Zhang X, Wu C, He Y, Ma Y, Hou H, Guo X, Du W, Zhao Y, Xia L: Engineering herbicide-resistant rice plants through CRISPR/Cas9-mediated homologous recombination of acetolactate synthase. Mol Plant 2016, 9:628–631.

7. Ren B, Yan F, Kuang Y, Li N, Zhang D, Zhou X, Lin H, Zhou H: Improved base editor for efficiently inducing genetic variations in rice with CRISPR/Cas9-guided hyperactive hAID mutant. Mol Plant 2018, 11:623–626.

8. Ren B, Yan F, Kuang Y, Li N, Zhang D, Lin H, Zhou H: A CRISPR/Cas9 toolkit for efficient targeted base editing to induce genetic variations in rice. Sci China Life Sci 2017, 60:516–519.

9. Yan F, Kuang Y, Ren B, Wang J, Zhang D, Lin H, Yang B, Zhou X, Zhou H: Highly efficient A.T to G.C base editing by Cas9n-guided tRNA adenosine deaminase in rice. Mol Plant 2018, 11:631–634.

10. Li C, Zong Y, Wang Y, Jin S, Zhang D, Song Q, Zhang R, Gao C: Expanded base editing in rice and wheat using a Cas9-adenosine deaminase fusion. Genome Biol 2018, 19:59.

11. Li C, Zong Y, Jin S, Zhu H, Lin D, Li S, Qiu J-L, Wang Y, Gao C: SWISS: multiplexed orthogonal genome editing in plants with a Cas9 nickase and engineered CRISPR RNA scaffolds. Genome Biol 2020, 21:141.

12. Zhao D, Li J, Li S, Xin X, Hu M, Price MA, Rosser SJ, Bi C, Zhang X: Glycosylase base editors enable C-to-A and C-to-G base changes. Nat Biotechnol 2020, DOI: 10.1038/s41587-020-0592-2.

13. Kurt IC, Zhou R, Iyer S, Garcia SP, Miller BR, Langner LM, Grunewald J, Joung JK: CRISPR C-to-G base editors for inducing targeted DNA transversions in human cells. Nat Biotechnol 2020, DOI: 10.1038/s41587-020-0609-x.

14. Lin Q, Zong Y, Xue C, Wang S, Jin S, Zhu Z, Wang Y, Anzalone AV, Raguram A, Doman JL, et al: Prime genome editing in rice and wheat. Nat Biotechnol 2020, 38:582–585.

15. Li H, Li J, Chen J, Yan L, Xia L: Precise modifications of both exogenous and endogenous genes in rice by prime editing. Mol Plant 2020, 13:671–674.

16. Tang X, Sretenovic S, Ren Q, Jia X, Li M, Fan T, Yin D, Xiang S, Guo Y, Liu L, et al: Plant prime editors enable precise gene editing in rice cells. Mol Plant 2020, 13:667–670.

17. Xu R, Li J, Liu X, Shan T, Qin R, Wei P: Development of plant prime-editing systems for precise genome editing. Plant Commun 2020, 1:100043.

18. Hsu PD, Scott DA, Weinstein JA, Ran FA, Konermann S, Agarwala V, Li Y, Fine EJ, Wu X, Shalem O, et al: DNA targeting specificity of RNA-guided Cas9 nucleases. Nat Biotechnol 2013, 31:827–832.

19. Sternberg SH, Redding S, Jinek M, Greene EC, Doudna JA: DNA interrogation by the CRISPR RNA-guided endonuclease Cas9. Nature 2014, 507:62–67.

20. Li Z, Liu ZB, Xing A, Moon BP, Koellhoffer JP, Huang L, Ward RT, Clifton E, Falco SC, Cigan AM: Cas9-guide RNA directed genome editing in soybean. Plant Physiol 2015, 169:960–970.

21. Lawrenson T, Shorinola O, Stacey N, Li CD, Ostergaard L, Patron N, Uauy C, Harwood W: Induction of targeted, heritable mutations in barley and *Brassica oleracea* using RNA-guided Cas9 nuclease. Genome Biology 2015, 16:258.

22. Barone P, Wu E, Lenderts B, Anand A, Gordon-Kamm W, Svitashev S, Kumar S: Efficient gene targeting in maize using inducible CRISPR-Cas9 and marker-free donor template. Mol Plant 2020,13:1219–1227.

23. Gosavi G, Yan F, Ren B, Kuang Y, Yan D, Zhou X, Zhou H: Applications of CRISPR technology in studying plant-pathogen interactions: overview and perspective. Phytopathol Res 2020, 2:21.

24. Kuang Y, Li S, Ren B, Yan F, Spetz C, Li X, Zhou X, Zhou H: Base-editing-mediated artificial evolution of *OsALS1 in planta* to develop novel herbicide-tolerant rice germplasms. Mol Plant 2020, 13:565–572.

25. Kleinstiver BP, Prew MS, Tsai SQ, Topkar VV, Nguyen NT, Zheng Z, Gonzales AP, Li Z, Peterson RT, Yeh JR, et al: Engineered CRISPR-Cas9 nucleases with altered PAM specificities. Nature 2015, 523:481–485.

26. Hu JH, Miller SM, Geurts MH, Tang W, Chen L, Sun N, Zeina CM, Gao X, Rees HA, Lin Z, Liu DR: Evolved Cas9 variants with broad PAM compatibility and high DNA specificity. Nature 2018, 556:57–63.

27. Nishimasu H, Shi X, Ishiguro S, Gao L, Hirano S, Okazaki S, Noda T, Abudayyeh OO, Gootenberg JS, Mori H, et al: Engineered CRISPR-Cas9 nuclease with expanded targeting space. Science 2018, 361:1259–1262.

28. Ren B, Liu L, Li S, Kuang Y, Wang J, Zhang D, Zhou X, Lin H, Zhou H: Cas9-NG greatly expands the targeting scope of the genome-editing toolkit by recognizing NG and other atypical PAMs in rice. Mol Plant 2019, 12:1015–1026.

29. Wang M, Xu Z, Gosavi G, Ren B, Cao Y, Kuang Y, Zhou C, Spetz C, Yan F, Zhou X, Zhou H: Targeted base editing in rice with CRISPR/ScCas9 system. Plant Biotechnol J 2020, 18:1645–1647.

30. Hua K, Tao X, Zhu JK: Expanding the base editing scope in rice by using Cas9 variants. Plant Biotechnol J 2019, 17:499–504.

31. Steinert J, Schiml S, Fauser F, Puchta H: Highly efficient heritable plant genome engineering using Cas9 orthologues from *Streptococcus thermophilus* and *Staphylococcus aureus*. Plant J 2015, 84:1295–1305.

32. Karvelis T, Gasiunas G, Young J, Bigelyte G, Silanskas A, Cigan M, Siksnys V: Rapid characterization of CRISPR-Cas9 protospacer adjacent motif sequence elements. Genome Biol 2015, 16:253.

33. Tang X, Lowder LG, Zhang T, Malzahn AA, Zheng X, Voytas DF, Zhong Z, Chen Y, Ren Q, Li Q, et al: A CRISPR-Cpf1 system for efficient genome editing and transcriptional repression in plants. Nat Plants 2017, 3:17103.

34. Ming M, Ren Q, Pan C, He Y, Zhang Y, Liu S, Zhong Z, Wang J, Malzahn AA, Wu J, et al: CRISPR–Cas12b enables efficient plant genome engineering. Nat Plants 2020, 6:202–208.

35. Walton RT, Christie KA, Whittaker MN, Kleinstiver BP: Unconstrained genome targeting with near-PAMless engineered CRISPR-Cas9 variants. Science 2020, 368:290–296.

36. Richter MF, Zhao KT, Eton E, Lapinaite A, Newby GA, Thuronyi BW, Wilson C, Koblan LW, Zeng J, Bauer DE, et al: Phage-assisted evolution of an adenine base editor with improved Cas domain compatibility and activity. Nat Biotechnol 2020, 38:883–891.

37. Lapinaite A, Knott GJ, Palumbo CM, Lin-Shiao E, Richter MF, Zhao KT, Beal PA, Liu DR, Doudna JA: DNA capture by a CRISPR-Cas9–guided adenine base editor. Science 2020, 369:566–571.

38. Hua K, Tao XP, Zhu JK: Expanding the base editing scope in rice by using Cas9 variants. Plant Biotechnol J 2019, 17:499–504.

39. Qin R, Li J, Liu X, Xu R, Yang J, Wei P: SpCas9-NG self-targets the sgRNA sequence in plant genome editing. Nat Plants 2020, 6:197–201.

40. Porebski S, Bailey LG, Baum BR: Modification of a CTAB DNA extraction protocol for plants containing high polysaccharide and polyphenol components. Plant Mol Biol Report 1997, 15:8–15.

