## Supplemental Figure1-18, and Supplemental Table1-3 for "SpRY greatly expands the genome editing scope in rice with highly flexible PAM recognition"

**Running title**: CRISPR/SpRY-mediated genome editing in rice

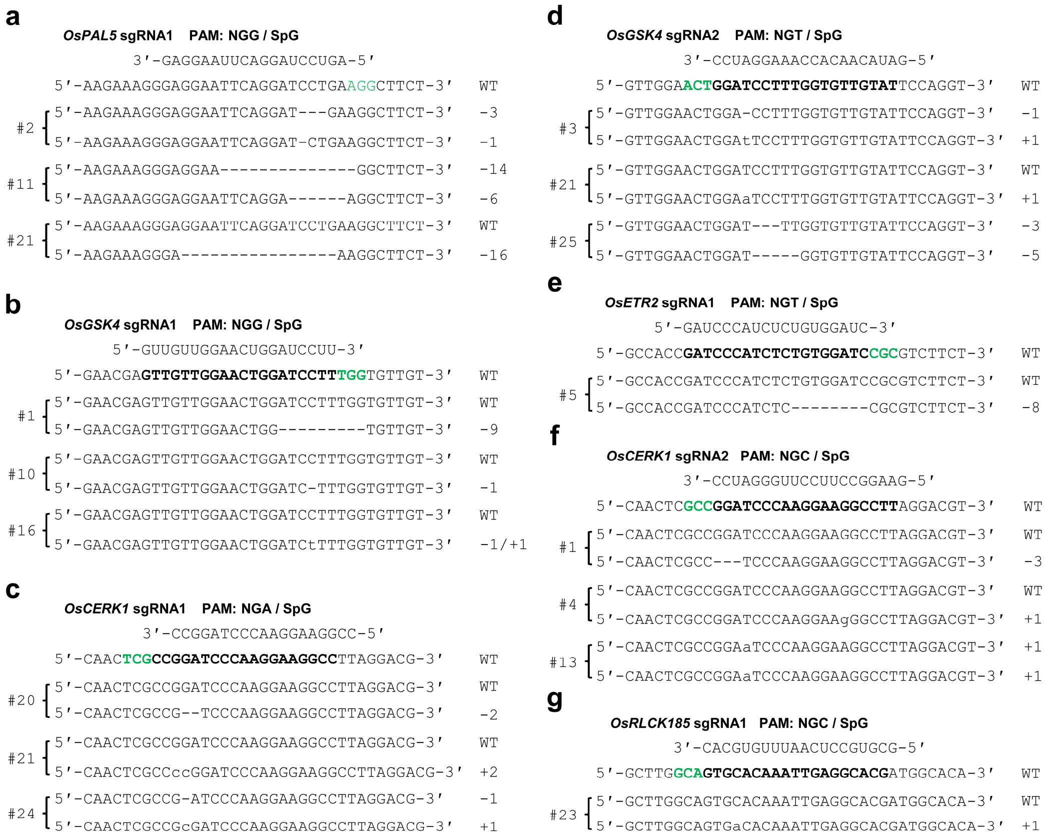


**Figure S1.** Analysis of SpG nuclease activity on different NGN PAMs in transgenic rice. **a-g** Representative mutant alleles of *OsPAL5* (**a**) and *OsGSK4* (**b**) with NGG PAMs, *OsCERK1* with an NGA PAM (**c**), *OsGSK4* (**d**) and *OsETR2* (**e**) with NGT PAMs, *OsCERK1* (**f**) and *OsRLCK185* (**g**) with an NGC PAM edited by SpG nuclease in T0 transgenic rice callus lines. WT, wild type; The PAM sequences and target sequences are highlighted in green and bold, respectively; nucleotide deletions and insertions are indicated by dashes and lowercase letters, respectively.

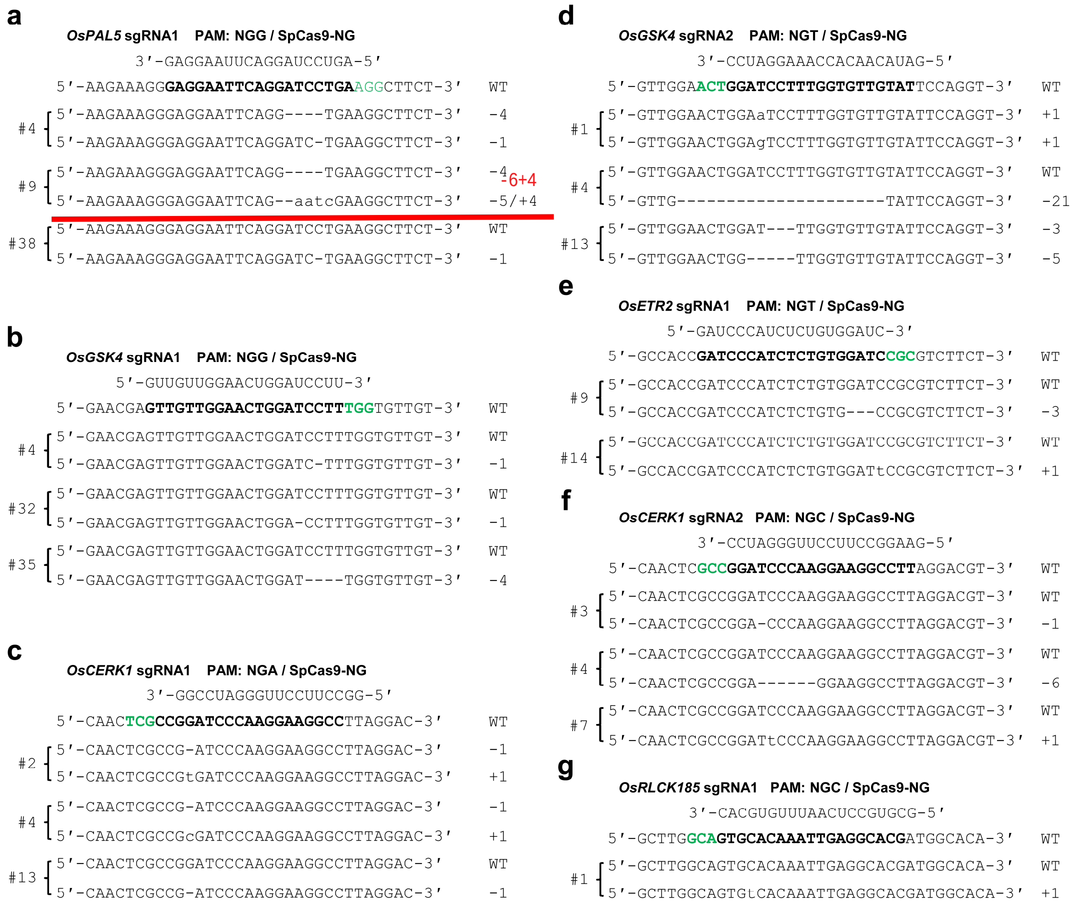

**Figure S2.** Analysis of SpCas9-NG nuclease activity on different NGN PAMs in transgenic rice. **a-g** Representative mutant alleles of *OsPAL5* (**a**) and *OsGSK4* (**b**) with NGG PAMs, *OsCERK1* with an NGA PAM (**c**), *OsGSK4* (**d**) and *OsETR2* (**e**) with NGT PAMs, *OsCERK1* (**f**) and *OsRLCK185* (**g**) with an NGC PAM edited by SpCas9-NG nuclease in T0 transgenic rice callus lines. WT, wild type; The PAM sequences and target sequences are highlighted in green and bold, respectively; nucleotide deletions and insertions are indicated by dashes and lowercase letters, respectively.

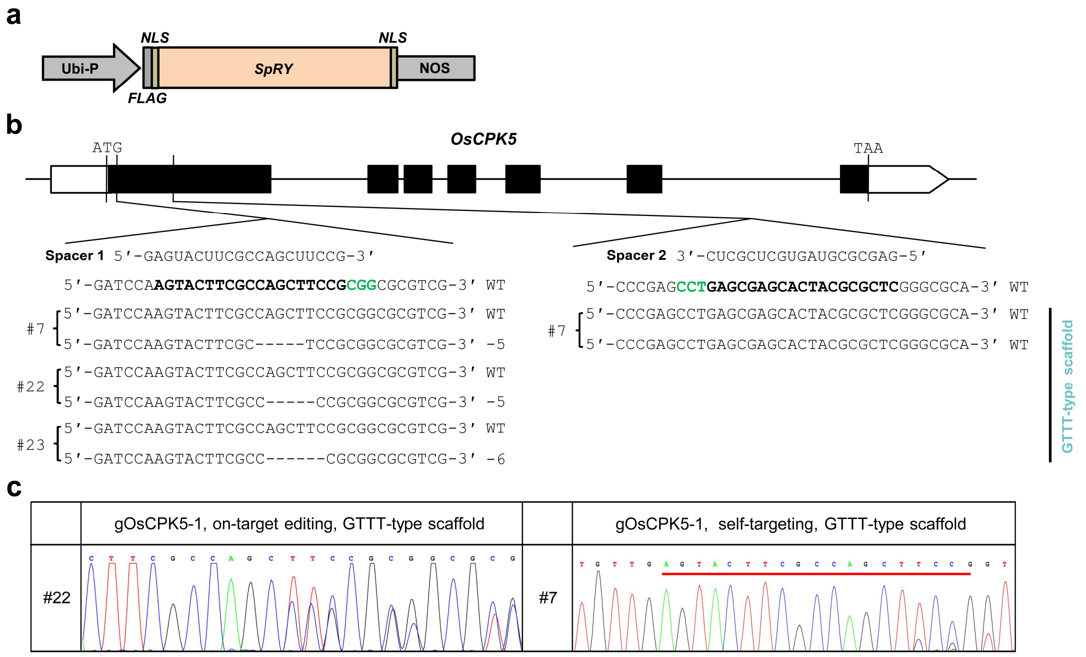


**Figure S3.** Targeted genome editing of *OsCPK5* by SpRY endonuclease in transgenic rice. **a** The gene construct of SpRY nuclease used for genome editing in transgenic rice. Ubi-P, maize ubiquitin 1 promoter; *NLS*, nuclear localization sequence. **b** Sequence results of the SpRY-edited *OsCPK5* at the NGG PAM sites using GTTT-type *sgRNA* scaffold in T0 transgenic rice callus lines. The exons are indicated by the black boxes; The PAM sequences and target sequences are highlighted in green and bold, respectively; Nucleotide deletions and insertions are indicated by dashes and lowercase letters, respectively. **c** Representative sanger sequencing chromatograms of the mutant *OsCPK5* allele and *OsCPK5-sgRNA* transgene in independent transgenic lines. The target region is underlined.

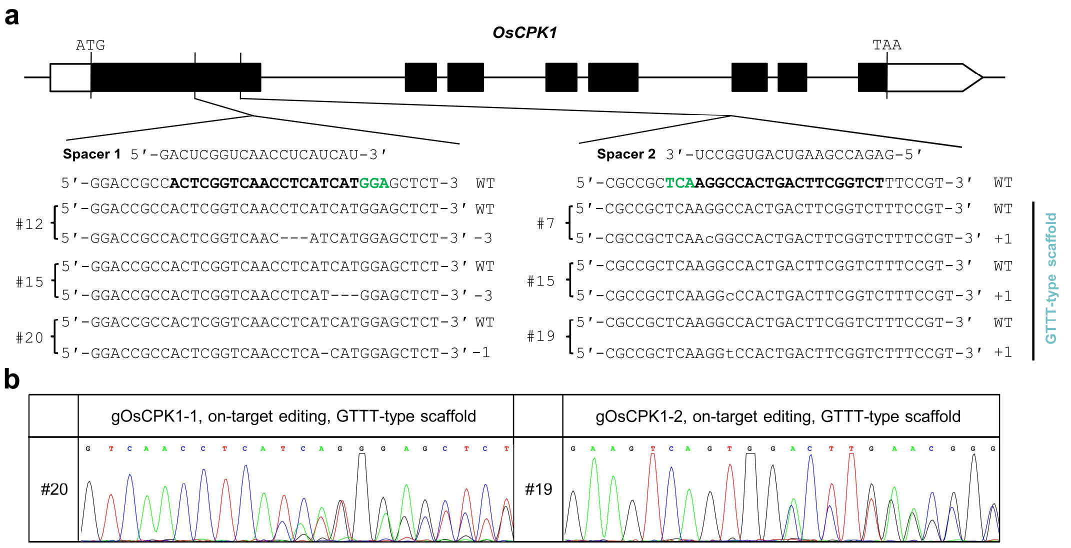


**Figure S4.** Targeted genome editing of *OsCPK1* by SpRY endonuclease in transgenic rice. **a** Sequence results of the SpRY-edited *OsCPK1* at the NGA PAM sites using GTTT-type *sgRNA* scaffold in T0 transgenic rice callus lines. The exons are indicated by the black boxes; The PAM sequences and target sequences are highlighted in green and bold, respectively; Nucleotide deletions and insertions are indicated by dashes and lowercase letters, respectively. **b** Representative sanger sequencing chromatograms of the mutant *OsCPK1* alleles in independent transgenic lines.

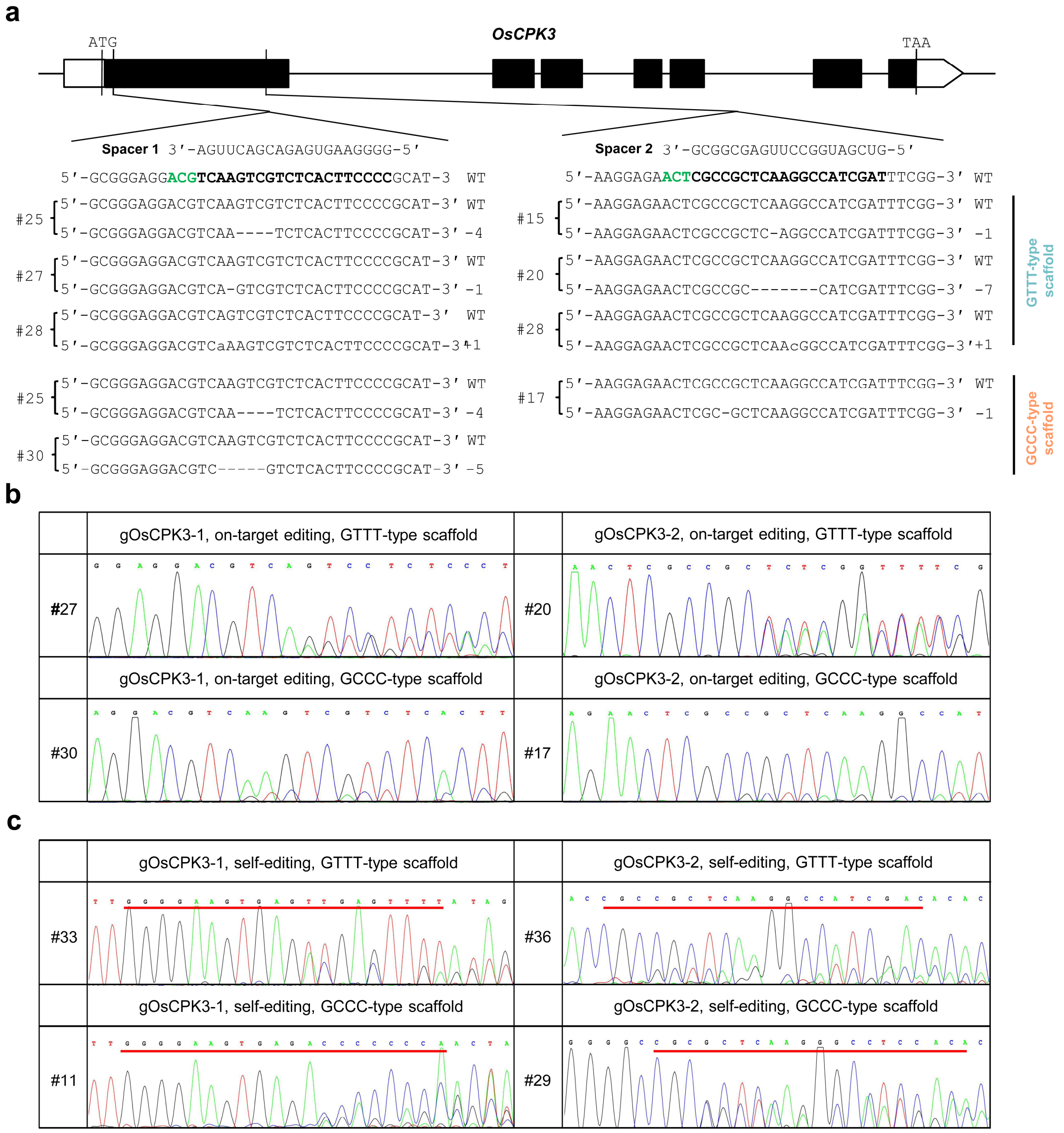


**Figure S5.** Targeted genome editing of *OsCPK3* by SpRY endonuclease in transgenic rice. **a** Sequence results of the SpRY-edited *OsCPK3* at the NGT PAM sites using GTTT-type and GCCC-type *sgRNA* scaffolds in T0 transgenic rice callus lines. The exons are indicated by the black boxes; The PAM sequences and target sequences are highlighted in green and bold, respectively; Nucleotide deletions and insertions are indicated by dashes and lowercase letters, respectively. **b** Representative sanger sequencing chromatograms of the mutant *OsCPK3* alleles in independent transgenic lines. **c** Representative sanger sequencing chromatograms of the mutant *OsCPK3-sgRNA* transgenes in independent transgenic lines. The target regions are underlined.

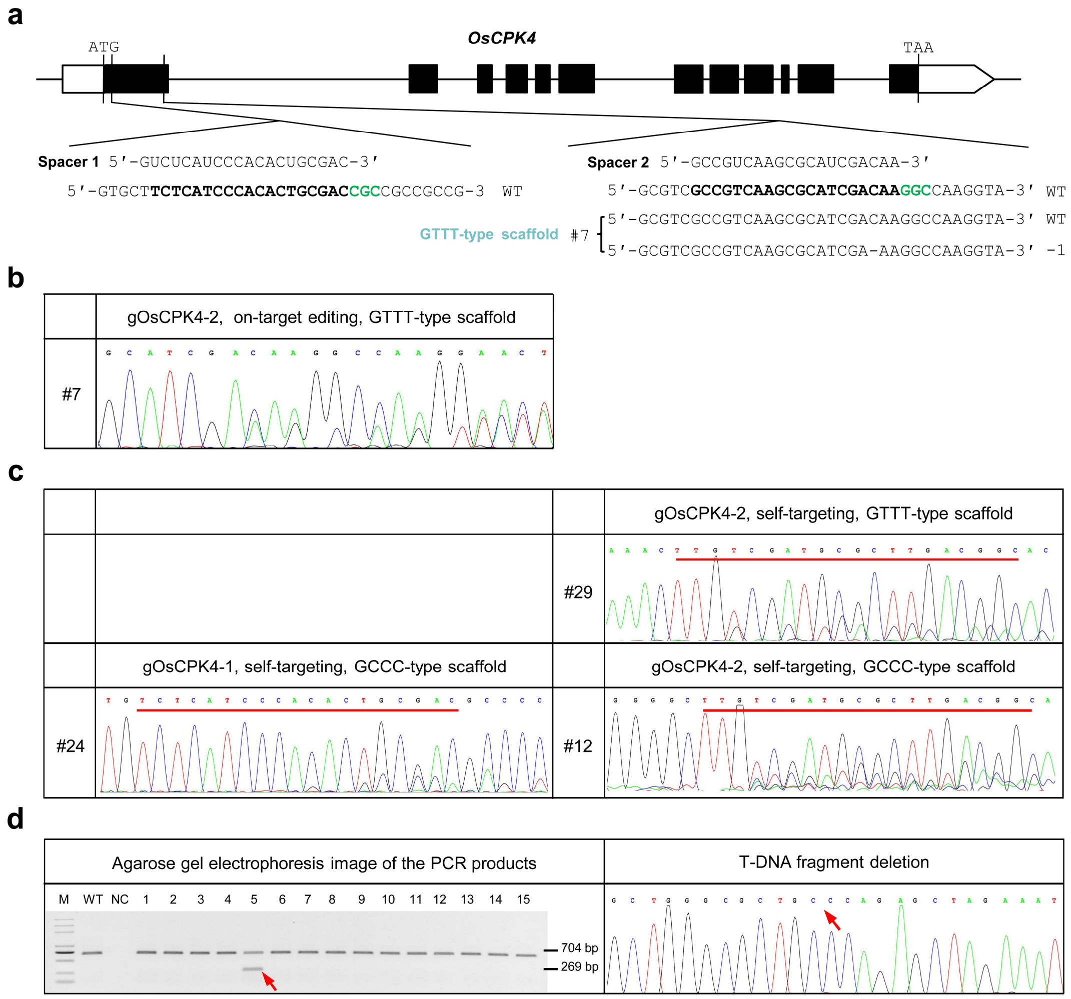


**Figure S6.** Targeted genome editing of *OsCPK4* by SpRY endonuclease in transgenic rice. **a** Sequence results of the SpRY-edited *OsCPK4* at the NGC PAM sites using GTTT-type *sgRNA* scaffold in T0 transgenic rice callus lines. The exons are indicated by the black boxes; The PAM sequences and target sequences are highlighted in green and bold, respectively; Nucleotide deletions and insertions are indicated by dashes and lowercase letters, respectively. **b** Representative sanger sequencing chromatogram of the mutant *OsCPK4* allele in independent transgenic line. **c** Representative sanger sequencing chromatograms of the mutant *OsCPK4-sgRNA* transgenes in independent transgenic callus lines. The target regions are underlined. **d** DNA fragment deletion between two *OsCPK4-sgRNA* transgenes induced by SpRY in transgenic lines. WT, wild type; NC, negative control. The deletion-junction site is marked by a red arrow.

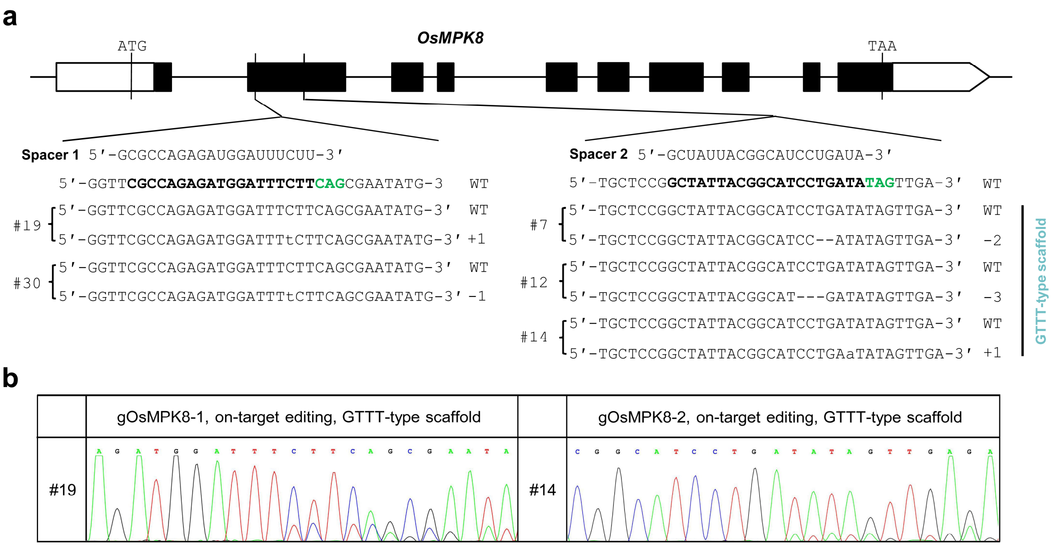


**Figure S7.** Targeted genome editing of *OsMPK8* by SpRY endonuclease in transgenic rice. **a** Sequence results of the SpRY-edited *OsMPK8* at the NAG PAM sites using GTTT-type *sgRNA* scaffold in T0 transgenic rice callus lines. The exons are indicated by the black boxes; The PAM sequences and target sequences are highlighted in green and bold, respectively; Nucleotide deletions and insertions are indicated by dashes and lowercase letters, respectively. **b** Representative sanger sequencing chromatograms of the mutant *OsMPK8* alleles in independent transgenic lines.

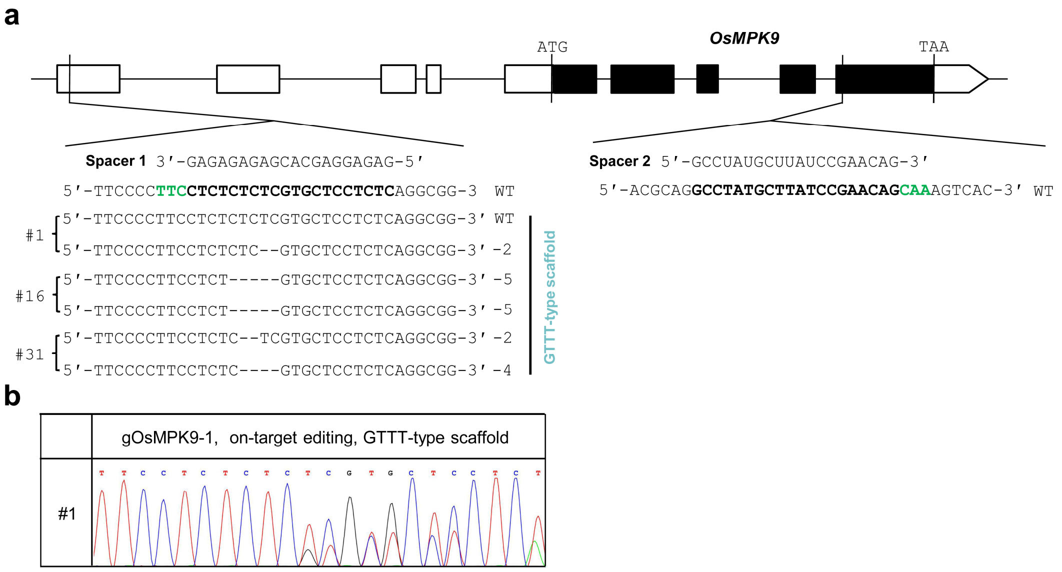


**Figure S8.** Targeted genome editing of *OsMPK9* by SpRY endonuclease in transgenic rice. **a** Sequence results of the SpRY-edited *OsMPK9* at the NAA PAM sites using GTTT-type *sgRNA* scaffold in T0 transgenic rice callus lines. The exons are indicated by the black boxes; The PAM sequences and target sequences are highlighted in green and bold, respectively; Nucleotide deletions and insertions are indicated by dashes and lowercase letters, respectively. **b** Representative sanger sequencing chromatogram of mutant *OsMPK9* allele in independent transgenic line.

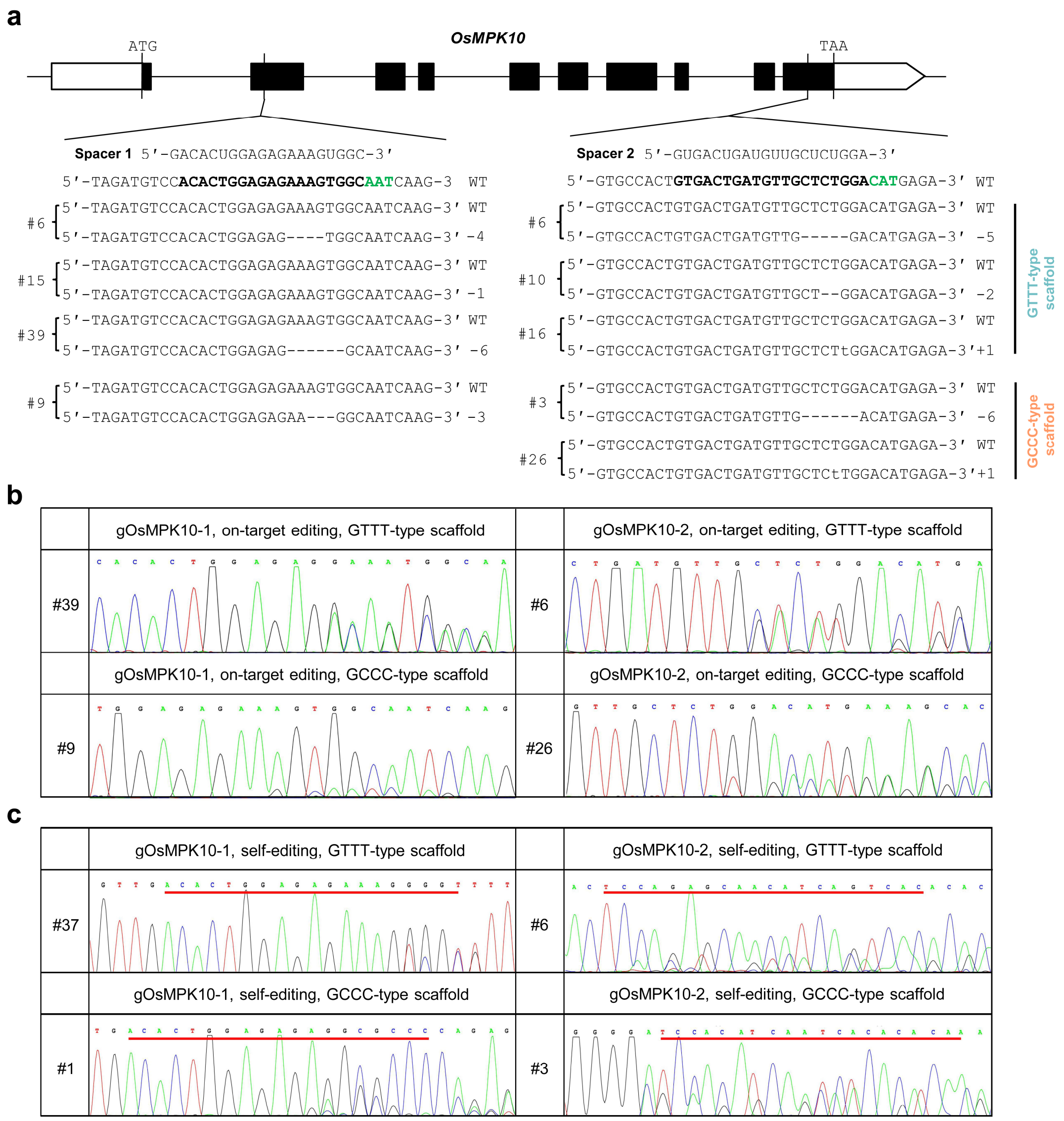


**Figure S9.** Targeted genome editing of *OsMPK10* by SpRY endonuclease in transgenic rice. **a** Sequence results of the SpRY-edited *OsMPK10* at the NAT PAM sites using GTTT-type and GCCC-type sgRNA scaffolds in T0 transgenic rice callus lines. The exons are indicated by the black boxes; The PAM sequences and target sequences are highlighted in green and bold, respectively; Nucleotide deletions and insertions are indicated by dashes and lowercase letters, respectively. **b** Representative sanger sequencing chromatograms of the mutant *OsMPK10* alleles in independent transgenic lines. **c** Representative sanger sequencing chromatograms of the mutant *OsMPK10-sgRNA* transgenes in independent transgenic lines. The target regions are underlined.

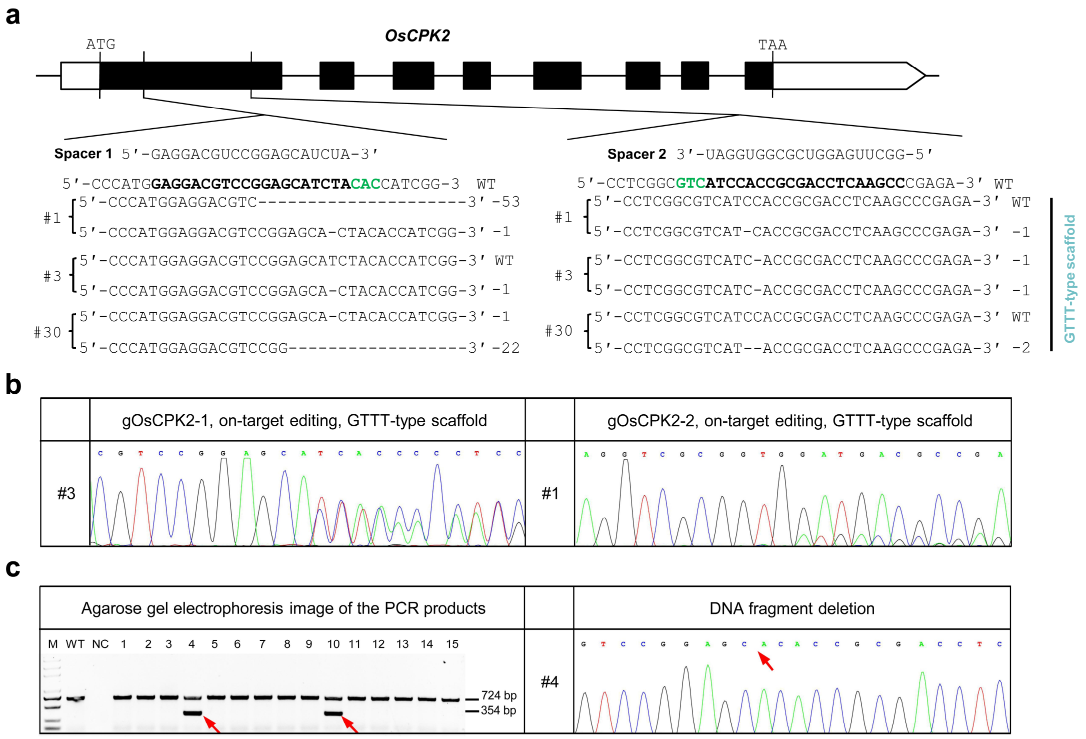


**Figure S10.** Targeted genome editing of *OsCPK2* by SpRY endonuclease in transgenic rice. **a** Sequence results of the SpRY-edited *OsCPK2* at the NAC PAM sites using GTTT-type sgRNA scaffold in T0 transgenic rice callus lines. The exons are indicated by the black boxes; The PAM sequences and target sequences are highlighted in green and bold, respectively; Nucleotide deletions and insertions are indicated by dashes and lowercase letters, respectively. **b** Representative sanger sequencing chromatograms of mutant *OsCPK2* alleles in independent transgenic lines. **c** DNA fragment deletion in *OsCPK2* induced by SpRY in transgenic line. WT, wild type; NC, negative control. The deletion-junction site is marked by a red arrow.

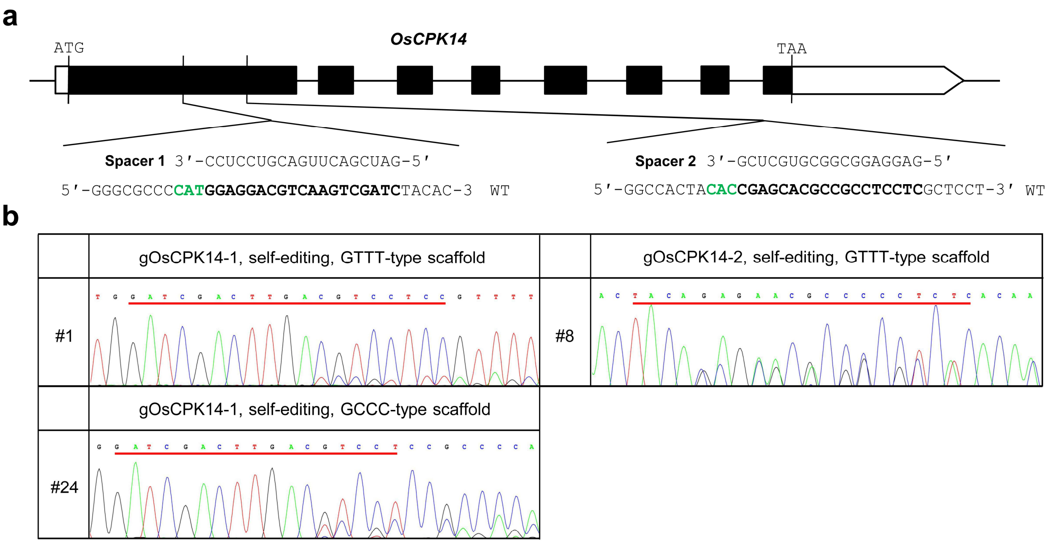


**Figure S11.** Targeted genome editing of *OsCPK14* by SpRY endonuclease in transgenic rice. **a** The target sites of *OsCPK14* in rice. The exons are indicated by the black boxes; The PAM sequences and target sequences are highlighted in green and bold, respectively. **b** Representative sanger sequencing chromatograms of the mutant *OsCPK14-sgRNA* transgenes in independent transgenic lines. The target regions are underlined.

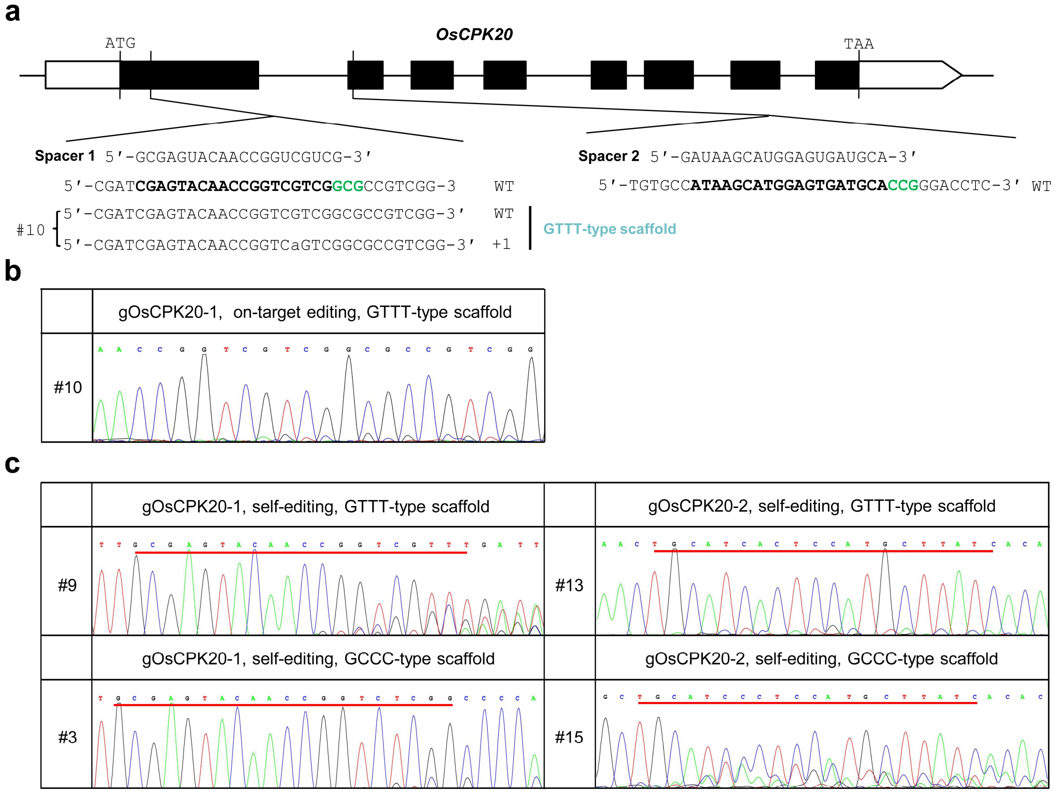


**Figure S12.** Targeted genome editing of *OsCPK20* by SpRY endonuclease in transgenic rice. **a** Sequence results of the SpRY-edited *OsCPK20* at the NCG PAM sites using GTTT-type *sgRNA* scaffold in T0 transgenic rice callus lines. The exons are indicated by the black boxes; The PAM sequences and target sequences are highlighted in green and bold, respectively; Nucleotide deletions and insertions are indicated by dashes and lowercase letters, respectively. **b** Representative sanger sequencing chromatogram of the mutant *OsCPK20* allele in independent transgenic line. **c** Representative sanger sequencing chromatograms of the mutant *OsCPK20-sgRNA* transgenes in independent transgenic lines. The target regions are underlined.

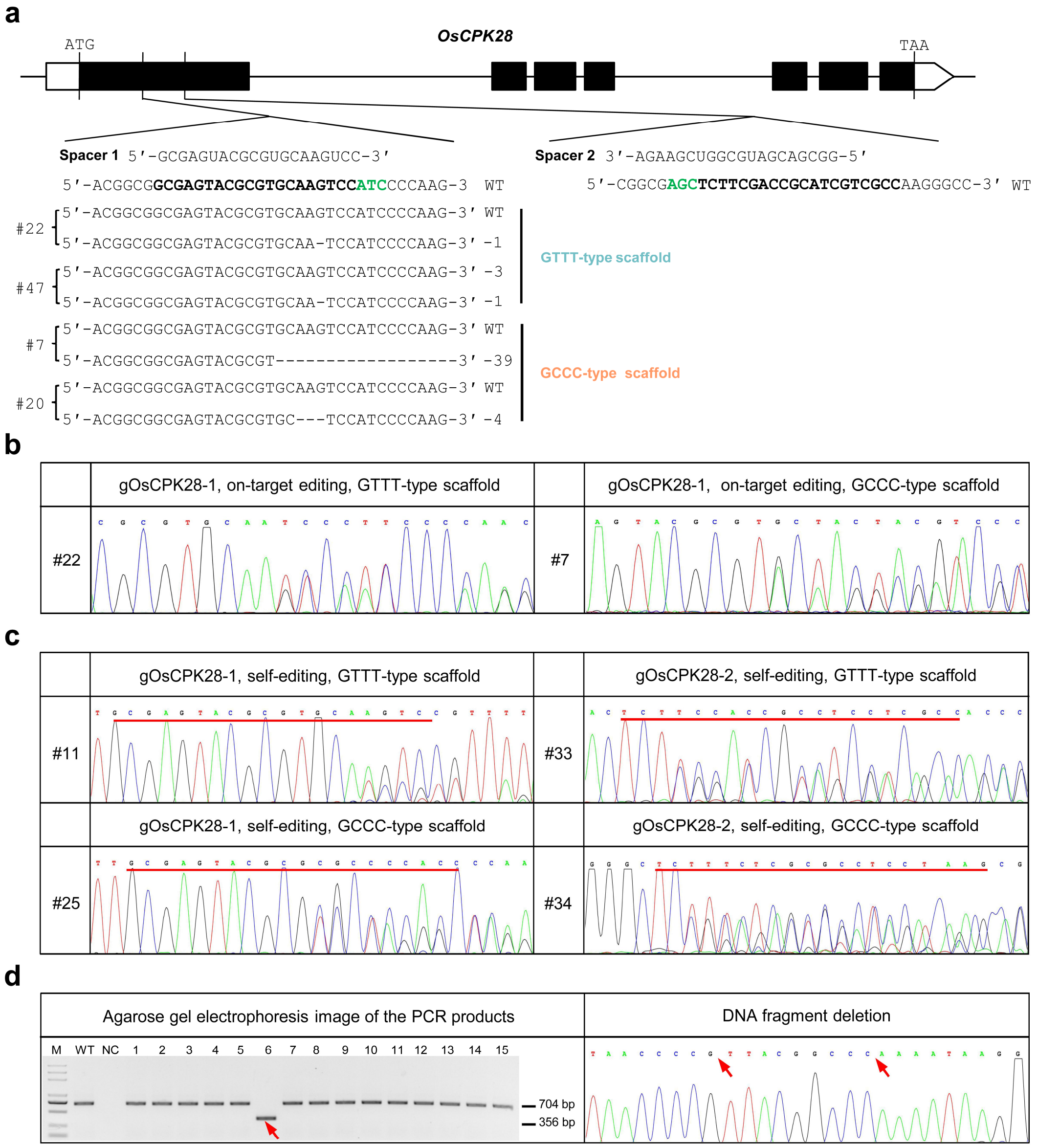


**Figure S13.** Targeted genome editing of *OsCPK28* by SpRY endonuclease in transgenic rice. **a** Sequence results of the SpRY-edited *OsCPK28* at the NTC and NCT PAM site using GTTT-type and GCCC-type sgRNA scaffolds in T0 transgenic rice callus lines. The exons are indicated by the black boxes; The PAM sequences and target sequences are highlighted in green and bold, respectively; Nucleotide deletions and insertions are indicated by dashes and lowercase letters, respectively. **b** Representative sanger sequencing chromatograms of the mutant *OsCPK28* alleles in independent transgenic lines. **c** Representative sanger sequencing chromatograms of the mutant *OsCPK28-sgRNA* transgenes in independent transgenic lines. The target regions are underlined. **d** DNA fragment deletion between two *OsCPK28-sgRNA* transgenes induced by SpRY in transgenic line. WT, wild type; NC, negative control. The deletion-junction site is marked by a red arrow.

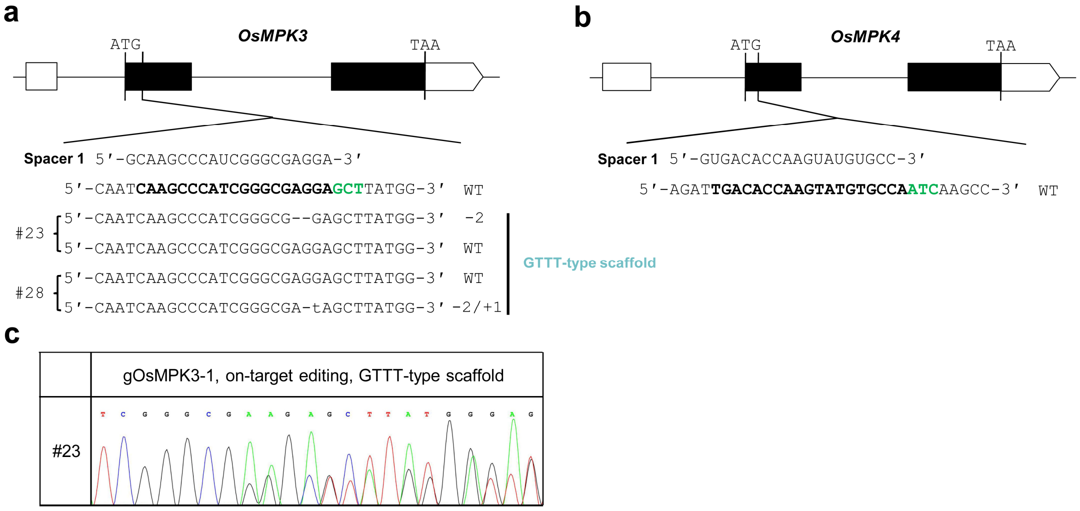


**Figure S14.** Targeted genome editing of *OsMPK3* and *OsMPK4* simultaneously by SpRY endonuclease in transgenic rice. **a** Sequence results of the SpRY-edited *OsMPK3* at the NCT PAM site using GTTT-type *sgRNA* scaffold in T0 transgenic rice callus lines. Nucleotide deletions and insertions are indicated by dashes and lowercase letters, respectively. **b** The target sites of *OsMPK4* in rice. In (**a** and **b**), the exons are indicated by the black boxes; The PAM sequences and target sequences are highlighted in green and bold, respectively; **c** Representative sanger sequencing chromatogram of mutant *OsMPK3* allele in transgenic line.

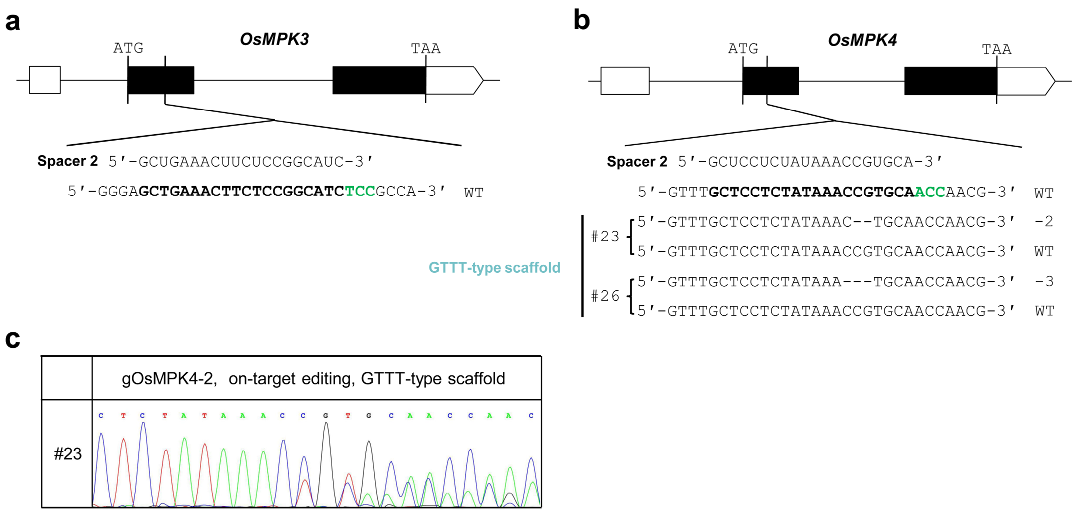


**Figure S15.** Targeted genome editing of *OsMPK3* and *OsMPK4* simultaneously by SpRY endonuclease in transgenic rice. **a** The target sites of *OsMPK3* in rice. **b** Sequence results of the SpRY-edited *OsMPK4* at the NCC PAM site using GTTT-type *sgRNA* scaffold in T0 transgenic rice callus lines. Nucleotide deletions and insertions are indicated by dashes and lowercase letters, respectively. In (**a** and **b**), the exons are indicated by the black boxes; The PAM sequences and target sequences are highlighted in green and bold, respectively; **c** Representative sanger sequencing chromatogram of mutant *OsMPK4* allele in transgenic line.

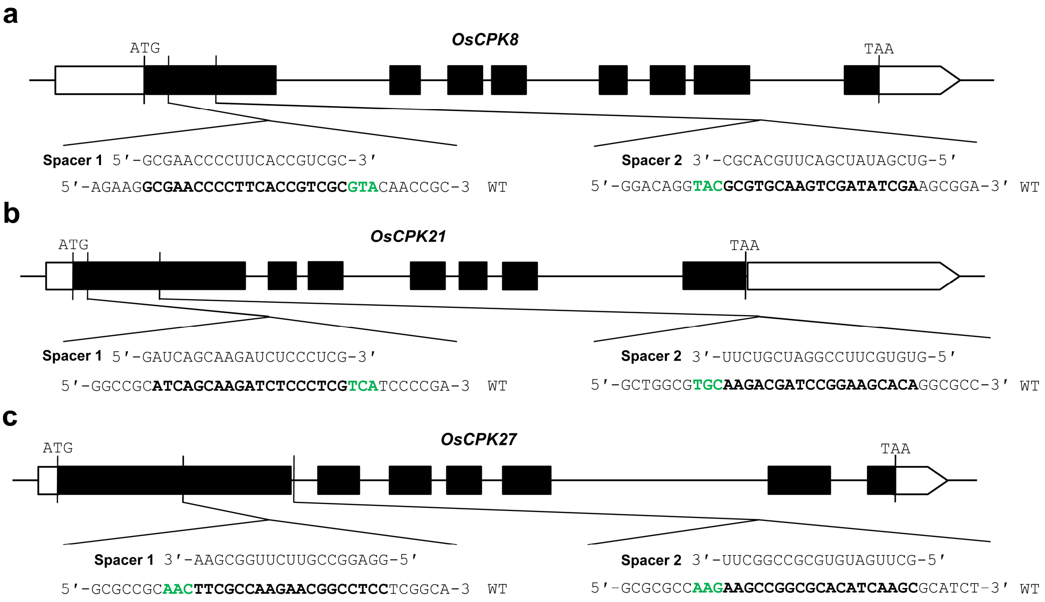


**Figure S16.** Targeted genome editing of *OsCPK8, OsCPK21,* and *OsCPK27* by SpRY endonuclease in transgenic rice. **a-c** The target sites of *OsCPK8* (**a**), *OsCPK21* (**b**), and *OsCPK27* (**c**) in rice. The exons are indicated by the black boxes; The PAM sequences and target sequences are highlighted in green and bold, respectively; Nucleotide deletions and insertions are indicated by dashes and lowercase letters, respectively.

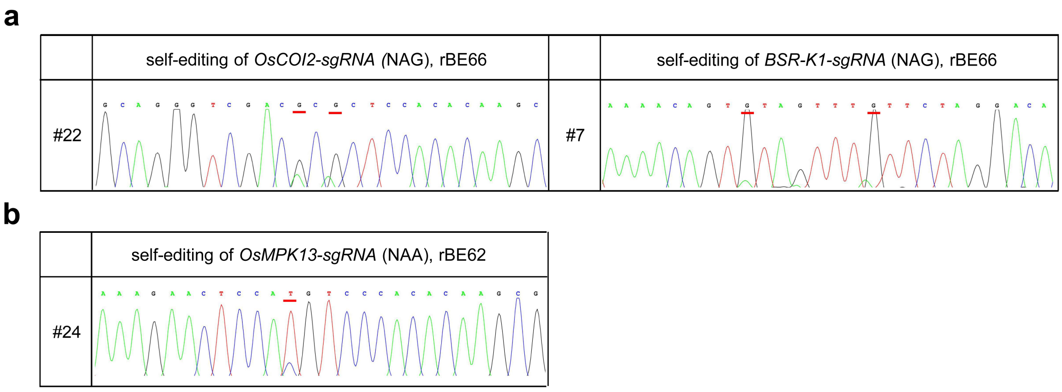


**Figure S17.** Self-editing activity of rBE66 and rBE62 in transgenic rice. **a** Representative sanger sequencing chromatograms of the *OsCOI2-sgRNA* and *BSR-K1-sgRNA* transgenes carrying nucleotide substitutions induced by rBE66 in independent transgenic callus lines. **b** Representative sanger sequencing chromatograms of the *OsMPK13-sgRNA* transgene carrying nucleotide substitution induced by rBE62 in independent transgenic callus line. In (**a** and **b**), the nucleotide substitutions are underlined.

**a**

***OsMPK8*-targeting sgRNA 1**

| | Target region | Gene | Freq. |
|---|---|---|---|
| On-target | CGCCAGAGATGGATTTCTTCAG | *Os01g0665200/OsMPK8* | 5.71% |
| Off-target 1 | CAACAGAGATGGATTTCTTCAG | *Os05g0566450* | 0.00% |

***OsMPK10*-targeting sgRNA 2**

| | Target region | Gene or Position | Freq. |
|---|---|---|---|
| On-target | GTGACTGATGTTGCTCTGGACAT | *Os01g0629900/OsMPK10* | 43.58% |
| Off-target 1 | ACGGCTGATGTTGCTCTGGACAT | *Os05g0582450* | 0.00% |
| Off-target 2 | CTGACTGATGTTGATCTGGTGGT | Chr12:10762404-2382 | 0.00% |
| Off-target 3 | GGGACAGATGTTGCTCTGGAATT | Chr08:17339974-9952 | 0.00% |

***OsCPK3*-targeting sgRNA 2**

| | Target region | Gene | Freq. |
|---|---|---|---|
| On-target | TCGATGGCCTTGAGCGGCGAGT | *Os01g0832300/OsCPK3* | 20.00% |
| Off-target 1 | TCGATGGCCTTGAGCGGCGCGT | *Os04g0560600/OsCPK12* | 0.00% |
| Off-target 2 | TCGATGACCTTGAGCGGCGAGT | *Os08g0540400/OsCPK21* | 15.00% |
| Off-target 3 | TCAGTGGCCTTGAGCGGCGAGT | *Os01g0622600/OsCPK1* | 0.00% |
| Off-target 4 | TCGATGGCCTTGAGCTGCGCGT | *Os03g0688300/OsCPK9* | 0.00% |
| Off-target 5 | TCGATGGTCTTGAGCGGCGCGT | *Os12g0486600/OsCPK27* | 0.00% |
| Off-target 6 | TCGGTGGCCTTGAGCGGCGCGT | *Os11g0136600/OsCPK25* | 2.50% |

***OsCPK28*-targeting sgRNA 1**

| | Target region | Gene | Freq. |
|---|---|---|---|
| On-target | GCGAGTACGCGTGCAAGTCCATC | *Os12g0169800/OsCPK28* | 73.91% |
| Off-target 1 | GCGAGTTCGCGTGCAAGTCCATC | *Os11g0171500/OsCPK24* | 0.00% |
| Off-target 2 | GCGAGTACGCGTGCAAGTCGATC | *Os12g0486600/OsCPK27* | 0.00% |

***OsCPK28*-targeting sgRNA 2**

| | Target region | Gene | Freq. |
|---|---|---|---|
| On target1 | AGCTCTTCGACCGCATCGTCGCC | *Os12g0169800/OsCPK28* | 0.00% |
| Off-target 1 | AGCTCTTCGACCGCATCGTCGCC | *Os11g0171500/OsCPK24* | 0.00% |
| Off-target 2 | AGCTCTTCGACCGCATCGTCGCC | *Os12g0230200/OsCPK29* | 6.52% |
| Off-target 3 | AGCTGTTCGACCGCATCGTCGCC | *Os05g0467150* | 0.00% |
| Off-target 4 | AGCTCTTCGACCGCATCATCGCC | *Os05g0585500/OsCPK15* | 0.00% |
| Off-target 5 | AGCTCTTCGACCGCATCTTCGCC | *Os09g0514200/OsCPK22* | 0.00% |
| Off-target 6 | AGCTCTTCGACCGGATCGTCGCC | *Os07g0568600/OsCPK20* | 0.00% |
| Off-target 7 | AGCTCTTCGACCGCATCGTCGCG | *Os01g0832300/OsCPK3* | 23.91% |
| Off-target 8 | AGCTCTTCGACCGCATCATCGCC | *Os01g0622600/OsCPK1* | 6.52% |

**b**

***OsCPK3*-targeting sgRNA 2 / off-target site 2: *OsCPK21***

5′-UCGAUGACCUUGAGCGGCG-3′

5′-GAAG**TCGATGACCTTGAGCGGCG**AGTCCTCCGATT-3′ WT

#30 5′-GAAGTCGATGACCTTGAGCGGCGAGTCCTCCGATT-3′ WT

5′-GAAGTCGATGACC-----CGGCGAGTCCTCCGATT-3′ -5

#46 5′-GAAGTCGATGACCTTGAGCGGCGAGTCCTCCGATT-3 WT

5′-GAAGTCGATGACCTTGAGCG-CGAGTCCTCCGATT-3 ′ -1

***OsCPK3*-targeting sgRNA 2 / off-target site 6: *OsCPK25***

3′-CGCCGCUCAAGGCCACCGA-5′

5′-GACGACG**CGCCGCTCAAGGCCACCGA**CTTCGGCCT-3 ′ WT

#34 5′-GACGACGCGCCGCTCAAGGCCACCGACTTCGGCCT-3 ′ WT

5′-GACGACGCG----TCAAGGCCACCGACTTCGGCCT-3 ′ -4

**c**

***OsCPK28*-targeting sgRNA 2 / off-target site: *OsCPK29***

3′-UCUUCGACCGCAUCGUCGCC-′

5′-GGCGAGC**TCTTCGACCGCATCGTCGCC**CGCGGCCA-3′ WT

#15 5′-GGCGAGCTCTTCGACCGCATCGTCGCCCGCGGCCA-3′ WT

5′-GGCGAGCT--------GCATCGTCGCCCGCGGCCA-3′ -8

#22 5′-GGCGAGCTCTT-----GCATCGTCGCCCGCGGCCA-3′ -5

5′-GGCG-----------CGCATCGTCGCCCGCGGCCA-3′ -11

***OsCPK28*-targeting sgRNA 7 / off-target site: *OsCPK3***

3′-UCUUCGACCGCAUCGUCGCG-5′

5′-GGGGAGC**TCTTCGACCGCATCGTCGCG**CGGGGCCA-3 ′ WT

#22 5′-GGGGAGCTCTTCGACCGCATCGTCGCGCGGGGCCA-3′ WT

5′-GGGGAGCTCT-----CGCATCGTCGCGCGGGGCCA-3 ′ -5

#27 5′-GGGGAGCTCTTCGACCGCATCGTCGCGCGGGGCCA-3 ′ WT

5′-GGGGAGCTCT-CGACCGCATCGTCGCGCGGGGCCA-3 ′ -1

#38 5′-GGGGAGCTCTTCGACCGCATCGTCGCGCGGGGCCA-3 ′ WT

5′-GGGGAGCTCTtTCGACCGCATCGTCGCGCGGGGCCA-3 ′ +T

#44 5′-GGGGAGCTCTTCGACCGCATCGTCGCGCGGGGCCA-3 ′ WT

5′-GGGGAGCTCTT--ACCGCATCGTCGCGCGGGGCCA-3 ′ -2

***OsCPK28*-targeting sgRNA 8 / off-target site: *OsCPK1***

3′-UCUUCGACCGCAUCAUCGCC-5′

5′-GGGGAGC**TCTTCGACCGCATCATCGCC**AGGGGGCA-3 ′ WT

#13 5′-GGGGAGCTCTTCGACCGCATCATCGCCAGGGGGCA-3 ′ WT

5′-GGGGAGCTC------CGCATCATCGCCAGGGGGCA-3 ′ -6

#35 5′-GGGGAGCTCTTCGACCGCATCATCGCCAGGGGGCA-3 ′ WT

5′-GGGGAGCTCT---ACCGCATCATCGCCAGGGGGCA-3 ′ -3

**Figure S18.** Off-target analysis of SpRY nuclease. **a** Summary of mutation frequencies in the potential off-target sites in the rice genome for SpRY nuclease in this study. The PAM sequences and the mismatches in the sgRNA sequences are highlighted in green and red, respectively. **b** Representative mutant alleles of off-targets detected in independent transgenic lines. WT, wild type; The PAM sequences and target sequences are highlighted in green and bold, respectively; nucleotide deletions and insertions are indicated by dashes and lowercase letters, respectively.

**Table S1. The complete nucleotide sequences of the rice codon-optimized *SpG*, *SpRY* and *sgRNA* fragments.**

| | |
|---|---|
| *SpG-fg1* | AAGCTTGGATCCGATCCAAATCCGAATCCGGCCATGGACTATAAGGATCACGATGGCGACTACAAGGATCATGACATTGACTATAAG<br>GATGACGACGATAAGATGGCACCTAAGAAGAAAAGGAAAGTCGGCATTCATGGCGTTCCGGCAGCCGACAAAAAGTATAGCATCGGC<br>CTCGATATTGGGACAAACTCTGTGGGCTGGGCGGTAATTACCGACGAGTACAAGGTGCCTAGTAAGAAATTTAAAGTGCTCGGAAAC<br>ACTGACAGGCACTCTATAAAGAAGAACCTGATCGGGGCACTGCTTTTCGACTCCGGAGAGACGGCGGAGGCGACGCGTCTCAAGCGT<br>ACCGCGCGCCGCAGGTACACAAGAAGGAAGAATAGGATCTGCTACTTGCAGGAAATCTTCAGTAACGAGATGGCGAAGGTCGACGAT<br>AGTTTCTTTCATCGGTTGGAAGAATCGTTCCTCGTAGAGGAGGACAAAAAGCACGAGCGTCACCCAATATTCGGGAATATTGTTGAC<br>GAGGTTGCCTACCATGAGAAATATCCTACAATATATCACCTCCGTAAGAAGCTTGTCGATTCAACTGATAAGGCTGATCTCAGACTC<br>ATCTATCTTGCCCTCGCACATATGATTAAGTTTCGTGGCCACTTCTTGATTGAAGGCGACCTCAACCCGGACAACTCAGATGTTGAC<br>AAGCTTTTTATACAGCTCGTCCAGACATATAACCAGCTGTTTGAAGAGAATCCCATCAATGCGAGTGGGGTTGATGCTAAAGCCATT<br>TTGTCCGCCAGGTTGTCCAAATCTCGCAGACTGGAAAACCTGATCGCACAGCTTCCCGGTGAAAAGAAAAACGGGCTCTTCGGCAAT<br>CTCATCGCACTGTCCCTCGGCCTCACCCCAAACTTCAAGTCTAACTTCGACCTGGCCGAGGATGCGAAGCTCCAGCTGTCAAAAGAT<br>ACATACGACGACGATTTGGACAATCTGCTTGCGCAAATAGGCGACCAGTATGCGGACCTGTTCCTGGCTGCCAAAAATCTGTCAGAT<br>GCAATCCTCCTGTCCGATATATTGCGTGTGAACACCGAAATCACGAAGGCACCGCTTAGCGCATCCATGATCAAGAGATACGACGAG<br>CACCATCAGGACCTCACACTCCTCAAGGCGCTTGTTCGTCAGCAGCTTCCCGAGAAATATAAGGAAATTTTTTTCGATCAAAGCAAG<br>AATGGATATGCTGGCTATATTGACGGTGGCGCTTCGCAGGAGGAGTTCTATAAATTCATTAAGCCGATTCTGGAGAAGATGGACGGA<br>ACGGAGGAGCTCCTCGTCAAGCTTAACCGGGAAGACCTGTTGCGGAAGCAGAGGACTTTTGATAACGGCTCTATTCCGCACCAAATC<br>CATCTGGGTGAGTTGCACGCAATCTTGAGAAGACAAGAGGATTTCTACCCGTTCCTTAAGGATAACAGAGAGAAGATAGAAAAAATA<br>CTGACCTTCAGGATACCATACTATGTGGGCCCACTGGCGCGCGGAAATAGTCGTTTCGCATGGATGACTAGAAAGTCCGAAGAAACG<br>ATCACGCCATGGAATTTTGAGGAAGTGGTCGACAAGGGCGCCTCTGCCCAGAGCTTCATCGAAAGGATGACCAATTTTGACAAAAAT<br>CTGCCTAACGAAAAGGTGCTTCCGAAGCACAGCCTGTTGTATGAATACTTCACAGTTTATAACGAGCTCACTAAGGTCAAGTACGTC<br>ACGGAGGGCATGCGTAAGCCTGCTTTCCTGTCTGGTGAACAAAAAAAGGCGATTGTGGACCTCCTTTTCAAGACGAACCGTAAAGTT<br>ACTGTGAAGCAACTGAAAGAGGATTACTTTAAGAAAATTGAGTGCTTCGACAGTGTGGAGATTTCCGGTGTCGAGGACCGGTTTAAC<br>GCCAGCCTGGGTACGTATCATGACCTGCTTAAAATTATCAAGGATAAAGATTTCCTGGATAATGAAGAGAACGAAGATATACTGGAG<br>GACATTGTGTTGACTTTGACCCTCTTCGAGGACAGAGAGATGATTGAGGAAAGACTGAAGACCTACGCACACCTTTTTGATGACAAG<br>GTCATGAAACAACTCAAGCGCCGGCGCTATACTGGCTGGGGCCGGCTTTCTCGCAAGCTCATCAATGGGATTCGGGATAAGCAATCA |

| | |
|---|---|
| | GGCAAGACAATTTTGGACTTCCTCAAATCCGACGGATTCGCAAATAGGAATTTTATGCAGCTGATACATGACGACTCTTTGACATTC<br>AAAGAAGACATACAGAAGGCTCAGGTCTCCGGCCAAGGAGATTCTTTGCACGAGCATATCGCTAACTTGGCAGGTAGCCCCGCCATA<br>AAAAAGGGCATTCTTCAAACGGTAAAAGTTGTTGACGAACTCGTGAAGGTTATGGGCCGTCATAAGCCGGAAAACATTGTTATTGAA<br>ATGGCTAGGGAAAATCAGACGACCCAGAAGGGACAGAAAAATAGCAGGGAGCGGATGAAGAGAATTGAAGAGGGAATTAAGGAGCTT<br>GGATCTCAGATTCTTAAGGAGCACCCTGTGGAGAACACCCAACTTCAGAATGAAAAGCTCTACCTTTACTACCTTCAAAACGGCCGG<br>GATATGTACGTCGATCAGGAACTTGACATTAACCGGTTGAGCGATTATGACGTTGACCATATTGTGCCCCAATCTTTCCTTAAAGAC<br>GACTCTATCGACAATAAAGTGCTGACGCGCAGCGATAAAAATCGCGGTAAGTCGGATAATGTCCCGTCGGAAGAGGTGGTTAAAAAA<br>ATGAAGAACTATTGGAGGCAACTCCTGAATGCCAAGCTGATCACTCAGAGGAAATTCGACAATCTCACCAAGGCAGAAAGGGGTGGA<br>CTTAGCGAGCTCGACAAGGCCGGTTTTATCAAAAGACAGCTGGTGGAGACACGCCAAATCACCAAACACGTTGCCCAGATCCTGGAT<br>TCGAGGATGAACACGAAGTATGACGAGAACGACAAGTTGATTAGGGAAGTCAAGGTCATCACTTTGAAGTCCAAGCTGGTGAGCGAC<br>TTTCGCAAAGACTTCCAGTTTTACAAAGTCAGGGAAATTAATAACTACCACCACGCCCACGACGCCTACCTTAACGCCGTGGTTGGC<br>ACAGCACTCATCAAGAAATACCCTAAGCTCGAATCTGAGTTCGTCTATGGCGACTATAAGGTCTACGACGTTAGAAAAATGATCGCG<br>AAATCTGAGCAGGAAATAGGCAAGGCAACTGCCAAGTACTTCTTCTATTCCAATATCATGAACTTTTTAAGACGGAGATTACCCTG<br>GCGAATGGTGAGATCCGCAAGCGCCCTTTGATTGAGA |
| *SpG-fg2* | AATGGTGAGATCCGCAAGCGCCCTTTGATTGAGACAAACGGAGAAACAGGAGAGATCGTATGGGACAAAGGGCGGGACTTTGCTACT<br>GTTAGGAAGGTGCTCTCTATGCCACAAGTTAACATTGTCAAAAAAACTGAAGTGCAGACAGGTGGGTTTAGCAAGGAATCTATCCTG<br>CCGAAGAGGAACTCTGACAAGCTGATCGCCCGCAAGAAAGATTGGGAcCCGAAAAAGTACGGAGGATTCTTGTGGCCCACAGTTGCG<br>TACTCCGTGCTTGTCGTGGCCAAAGTGGAGAAGGGCAAGTCTAAGAAGCTCAAGAGCGTCAAAGAGTTGTTGGGGATCACGATTATG<br>GAGCGGTCGTCTTTCGAAAAGAATCCGATAGATTTTCTCGAGGCCAAGGGTTATAAAGAAGTCAAGAAGGATCTTATCATCAAGCTC<br>CCTAAGTACTCCCTCTTTGAGCTTGAAAACGGACGGAAAAGAATGCTGGCTTCAGCGAAGCAGCTTCAGAAGGGTAATGAACTCGCT<br>CTGCCCTCAAAATATGTGAATTTCCTTTACCTGGCATCACACTATGAGAAGCTTAAGGGGTCTCCAGAGGACAACGAGCAGAAGCAA<br>CTGTTCGTTGAACAACACAAGCACTACCTTGACGAGATTATCGAGCAAATCAGCGAGTTTAGCAAGCGCGTTATACTGGCAGACGCA<br>AATCTTGATAAGGTCCTTAGCGCCTACAACAAGCATAGAGACAAACCCATCCGGGAGCAGGCCGAGAACATTATTCATCTCTTCACC<br>TTGACGAATCTTGGGGCCCCGGCCGCGTTCAAGTACTTCGATACTACCATAGACAGAAAGCAATATCGGTCGACAAAGGAAGTTCTT<br>GACGCCACGCTGATCCACCAAAGTATAACAGGCCTCTATGAGACACGCATCGACCTTTCGCAGTTGGGCGGTGACCGCCCCAAAAAG<br>AAGAGGAAAGTTGGCGGGTGAACTAGTGAATTC |

| | |
|---|---|
| *SpRY-fg2* | AATGGTGAGATCCGCAAGCGCCCTTTGATTGAGACAAACGGAGAAACAGGAGAGATCGTATGGGACAAAGGGCGGGACTTTGCTACT<br>GTTAGGAAGGTGCTCTCTATGCCACAAGTTAACATTGTCAAAAAAACTGAAGTGCAGACAGGTGGGTTTAGCAAGGAATCTATCAGG<br>CCGAAGAGGAACTCTGACAAGCTGATCGCCCGCAAGAAAGATTGGGAcCCGAAAAAGTACGGAGGATTCTTGTGGCCCACAGTTGCG<br>TACTCCGTGCTTGTCGTGGCCAAAGTGGAGAAGGGCAAGTCTAAGAAGCTCAAGAGCGTCAAAGAGTTGTTGGGGATCACGATTATG<br>GAGCGGTCGTCTTTCGAAAAGAATCCGATAGATTTTCTCGAGGCCAAGGGTTATAAAGAAGTCAAGAAGGATCTTATCATCAAGCTC<br>CCTAAGTACTCCCTCTTTGAGCTTGAAAACGGACGGAAAAGAATGCTGGCTTCAGCGAAGCAGCTTCAGAAGGGTAATGAACTCGCT<br>CTGCCCTCAAAATATGTGAATTTCCTTTACCTGGCATCACACTATGAGAAGCTTAAGGGGTCTCCAGAGGACAACGAGCAGAAGCAA<br>CTGTTCGTTGAACAACACAAGCACTACCTTGACGAGATTATCGAGCAAATCAGCGAGTTTAGCAAGCGCGTTATACTGGCAGACGCA<br>AATCTTGATAAGGTCCTTAGCGCCTACAACAAGCATAGAGACAAACCCATCCGGGAGCAGGCCGAGAACATTATTCATCTCTTCACC<br>TTGACGAGGCTTGGGGCCCCGAGAGCGTTCAAGTACTTCGATACTACCATAGACCCAAAGCAATATCGGTCGACAAAGGAAGTTCTT<br>GACGCCACGCTGATCCACCAAAGTATAACAGGCCTCTATGAGACACGCATCGACCTTTCGCAGTTGGGCGGTGACCGCCCCAAAAAG<br>AAGAGGAAAGTTGGCGGGTGAACTAGTGAATTC |
| GCCC-type *sgRNA* | GTCAGCGATGAGTACAGCAAGCCCCAGAGCTAGAAATAGCAAGTTGGGGTAAGGCTAGTCCGTTATCAACTTGAAAAAGTGGCACCG<br>AGTCGGTGCTTTTTTTTTGAGATTTCCAACCAGGTCCCTGGAGCCCATAGtctagtAACGGCCGCCAGTGTGCTGGAATTGCCCTTGG<br>ATCATGAACCAACGGCCTGGCTGTATTTGGTGGTTGTGTAGGGAGATGGGGAGAAGAAAAGCCCGATTCTCTTCGCTGTGATGGGCT<br>GGATGCATGCGGGGGAGCGGGAGGCCCAAGTACGTGCACGGTGAGCGGCCCACAGGGCGAGTGTGAGCGCGAGAGGCGGGAGGAACA<br>GTTTAGTACCACATTGCCCAGCTAACTCGAACGCGACCAACTTATAAACCCGCGCGCTGTCGCTTGTGTAGAGACCAAAGGAGGTCT<br>CAGCCCCAGAGCTAGAAATAGCAAGTTGGGGTAAGGCTAGTCCGTTATCAACTTGAAAAAGTGGCACCGAGTCGGTGCTTTTTTTGT<br>CCCTTCGAAGGGCAATTC |
| *TadA8e* | ATGTCAGAAGTCGAGTTCTCCCATGAGTATTGGATGAGGCACGCCCTCACTCTTGCGAAGAGGGCCAGGGACGAGAGGGAGGTGCCG<br>GTCGGTGCTGTCCTGGTCTTGAATAACAGGGTGATAGGCGAAGGTTGGAACAGGGCTATTGGCCTTCATGACCCTACTGCTCATGCG<br>GAAATCATGGCACTTAGACAGGGGGGCCTCGTTATGCAAAATTACCGCCTGATCGACGCCACTCTTTATGTCACATTTGAACCATGT<br>GTTATGTGTGCGGGCGCTATGATCCATTCACGCATAGGTCGCGTGGTTTTTGGAGTTCGCAACAGTAAACGTGGGGCTGCAGGCTCT<br>CTGATGAACGTTTTGAATTATCCGGGAATGAACCATAGAGTCGAAATCACAGAAGGGATTTTGGCAGACGAATGCGCGGCTCTTCTT<br>TGTGATTTTTACAGAATGCCCCGCCAAGTGTTTAATGCTCAAAAGAAAGCGCAGAGTAGCATCAACTCGGGGGGATCTTCTGGGGGC<br>TCGTCTGGTTCCGAGACTCCCGGAACTTCCGAGTCGGCAACACCTGAATCCTCCGGCGGCTCTTCGGGCGGATCTGAC |

| | |
|---|---|
| | AAAAAATACTCAATTGGTCTGGCTATTGGGACAAACTCTG |

**Table S2. List of oligonucleotides in this study.**

| Primer name | Primer sequence (5' - 3') | Used for |
|---|---|---|
| SpG-F3 | GCGAATGCATCTAGATATCGGATCCGATCCAAATCCGAA | Assembling the full-length *SpG* and *SpRY* genes through overlapping-extension PCR-based method |
| SpG-R3 | CTCTCCTGTTTCTCCGTTTGTCTCAATCAAAGGGCGCTTG | |
| SpG-F1a | CAAGCGCCCTTTGATTGAGACAAACGGAGAAACAGGAG | |
| SpG-R2 | TTCACTAGTTCACCCGCCAAC | |
| SpRY-F1 | AGGACGCGTCTCAAGCGTAC | Introducing A61R point mutation into SpG-fg1 |
| SpRY-R1 | CTCCGCCGTCTCTCCGGAGT | |
| UGI-F1 | TCCGGCGGAAGTACAAAC | Fusing *hAID*Δ* and *UGI* to *SpRY* at the 5'and 3' ends, resulting in *rBE66* |
| rAPO-R1 | AGCAAGTCCGATTGAATACT | |
| OsCas9-Fg1-F1 | ATTGGGACAAACTCTGTGG | |
| OsCas9-Fg2-R1 | GTCACCGCCCAACTGCGA | |
| TadA8e-F1 | CTTGGATCCATGTCAGAAGTCGAGTTC | Fusing *TadA8e* to the 5' end of *SpRY*, resulting in *rBE62* |
| TadA8e-F2 | CAGAGTTTGTCCCAATAGCC | |
| SpRY-F2 | GGCTATTGGGACAAACTCTG | |
| SpRY-R2 | TTCACTAGTTCACCCGCCAACTTTC | |
| gRNA4-NG-F1 | GTCCCTTCGAAGGGCAATTC | Generating pENTR4:sgRNA4-NG construct |
| gRNA4-NG-R1 | CTTGCTGTACTCATCGCTGAC | |

| | | |
|---|---|---|
| gRNA4-NG-F2 | GTCAGCGATGAGTACAGCAAG | |
| gRNA4-NG-R2 | GAATTGCCCTTCGAAGGGAC | |
| gOsCERK1-F3a | TGTTGGCCTTCCTTGGGATCCGG | Knocking out the endogenous *OsCERK1* gene with an NGA PAM |
| gOsCERK1-R3a | AAACCCGGATCCCAAGGAAGGCC | |
| OsCERK1-F3 | GACGTCTACGCCTTTGGTGT | Detecting nucleotide changes of *OsCERK1* |
| OsCERK1-R3 | GTCAGCTGCAAAATGCAATG | |
| gOsRLCK185-F1a | GTGTGCGTGCCTCAATTTGTGCAC | Knocking out the endogenous *OsRLCK185* gene with an NGC PAM |
| gOsRLCK185-R1a | AAACGTGCACAAATTGAGGCACG | |
| OsRLCK185-F2 | TCCATGGCCTTGTTCCTCTT | Detecting nucleotide changes of *OsRLCK185* |
| OsRLCK185-R2 | TGCTGCTAGACACATCCACA | |
| gOsGSK4-FGG | GTGTGTTGTTGGAACTGGATCCTT | Knocking out the endogenous *OsGSK4* gene with an NGG PAM |
| gOsGSK4-RGG | AAACAAGGATCCAGTTCCAACAAC | |
| OsGSK4-F1 | GGAAATGATCCAGTCACAGGT | Detecting nucleotide changes of *OsGSK4* |
| OsGSK4-R1 | TCGGGAACAAACTCCATGAC | |
| gOsPAL5-F2 | TGTTGGAGGAATTCAGGATCCTGA | Knocking out the endogenous *OsPAL5* gene with an NGG PAM |
| gOsPAL5-R2 | AAACTCAGGATCCTGAATTCCTCC | |
| OsPAL5-F1 | AACAACGGAATCAGAAAGCTG | Detecting nucleotide changes of *OsPAL5* |

| | | |
|---|---|---|
| OsPAL5-R1 | CACCGGAGAGAAGCTCAAGT | |
| gOsGSK4-FGT | TGTTGATACAACACCAAAGGATCC | Knocking out the endogenous *OsGSK4* gene with an NGT PAM |
| gOsGSK4-RGT | AAACGGATCCTTTGGTGTTGTATC | |
| OsGSK4-F1 | GGAAATGATCCAGTCACAGGT | Detecting nucleotide changes of *OsGSK4* |
| OsGSK4-R1 | TCGGGAACAAACTCCATGAC | |
| gOsETR2-FGT | GTGTGTCCCATCTCTGTGGATCCG | Knocking out the endogenous *OsETR2* gene with an NGT PAM |
| gOsETR2-RGT | AAACCGGATCCACAGAGATGGGAC | |
| OsETR2-F1 | TGGTTCGTTCGTTTGTTTGA | Detecting nucleotide changes of *OsETR2* |
| OsETR2-R1 | ATGGTGATGAGGTGGGTGAG | |
| gOsCERK1-FGC | TGTTGAAGGCCTTCCTTGGGATCC | Knocking out the endogenous *OsCERK1* gene with an NGC PAM |
| gOsCERK1-RGC | AAACGGATCCCAAGGAAGGCCTTC | |
| OsCERK1-F3 | GACGTCTACGCCTTTGGTGT | Detecting nucleotide changes of *OsCERK1* |
| OsCERK1-R3 | GTCAGCTGCAAAATGCAATG | |
| gOsMPK8-F2 | TGTTGCGCCAGAGATGGATTTCTT | Knocking out the endogenous *OsMPK8* gene using the GTTT-type sgRNA scaffold |
| gOsMPK8-R2 | AAACAAGAAATCCATCTCTGGCGC | |
| gOsMPK8-F3 | GTGTGCTATTACGGCATCCTGATA | Knocking out the endogenous *OsMPK8* gene using the GTTT-type sgRNA scaffold |
| gOsMPK8-R3 | AAACTATCAGGATGCCGTAATAGC | |

| | | |
|---|---|---|
| OsMPK8-F | CTTTGCATCAGAAGGGCAGG | Detecting nucleotide changes of *OsMPK8* |
| OsMPK8-R | CAGTCATCAATAAACGCTGTGCT | |
| gOsMPK9-F3 | TGTTGAGAGGAGCACGAGAGAGAG | Knocking out the endogenous *OsMPK9* gene using the GTTT-type sgRNA scaffold |
| gOsMPK9-R3 | AAACCTCTCTCTCGTGCTCCTCTC | |
| gOsMPK9-F4 | GTGTGCCTATGCTTATCCGAACAG | Knocking out the endogenous *OsMPK9* gene using the GTTT-type sgRNA scaffold |
| gOsMPK9-R4 | AAACCTGTTCGGATAAGCATAGGC | |
| OsMPK9-F3 | TTTACGGCACCACGAATCCA | Detecting nucleotide changes of *OsMPK9* |
| OsMPK9-R3 | AACATCAACGGGCACCCATA | |
| OsMPK9-F4 | CCAGGTGCACTTGTGTTTCA | |
| OsMPK9-R4 | TGTTAGTACCAACGCCTGCC | |
| gOsMPK10-F2 | TGTTGACACTGGAGAGAAAGTGGC | Knocking out the endogenous *OsMPK10* gene using the GTTT-type sgRNA scaffold |
| gOsMPK10-R2 | AAACGCCACTTTCTCTCCAGTGTC | |
| gOsMPK10-F3 | GTGTGTGACTGATGTTGCTCTGGA | Knocking out the endogenous *OsMPK10* gene using the GTTT-type sgRNA scaffold |
| gOsMPK10-R3 | AAACTCCAGAGCAACATCAGTCAC | |
| gOsMPK10-F2 | TGTTGACACTGGAGAGAAAGTGGC | Knocking out the endogenous *OsMPK10* gene using the GCCC-type sgRNA scaffold |
| gOsMPK10-NG-R2 | GGGCGCCACTTTCTCTCCAGTGTC | |
| gOsMPK10-F3 | GTGTGTGACTGATGTTGCTCTGGA | Knocking out the endogenous *OsMPK10* gene |

| | | |
|---|---|---|
| gOsMPK10-NG-R3 | GGGCTCCAGAGCAACATCAGTCAC | using the GCCC-type sgRNA scaffold |
| OsMPK10-F2 | TGATTGTGAGGACAGACGGT | Detecting nucleotide changes of *OsMPK10* |
| OsMPK10-R2 | GTGACAGTTCCCTACCAGTGT | |
| OsMPK10-F3 | CCAACAGGTGCTTTGTGGTTAAT | |
| OsMPK10-R3 | AGGGAACAACACCGACCTTT | |
| gOsCPK1-F1 | TGTTGACTCGGTCAACCTCATCAT | Knocking out the endogenous *OsCPK1* gene using the GTTT-type sgRNA scaffold |
| gOsCPK1-R1 | AAACATGATGAGGTTGACCGAGTC | |
| gOsCPK1-F2 | GTGTGAGACCGAAGTCAGTGGCCT | Knocking out the endogenous *OsCPK1* gene using the GTTT-type sgRNA scaffold |
| gOsCPK1-R2 | AAACAGGCCACTGACTTCGGTCTC | |
| OsCPK1-F | GTCACCTACCTCGTCACCCA | Detecting nucleotide changes of *OsCPK1* |
| OsCPK1-R | TTTCAGACCTGTCAAACCCA | |
| gOsCPK2-F1 | TGTTGAGGACGTCCGGAGCATCTA | Knocking out the endogenous *OsCPK2* gene using the GTTT-type sgRNA scaffold |
| gOsCPK2-R1 | AAACTAGATGCTCCGGACGTCCTC | |
| gOsCPK2-F2 | GTGTGGCTTGAGGTCGCGGTGGAT | Knocking out the endogenous *OsCPK2* gene using the GTTT-type sgRNA scaffold |
| gOsCPK2-R2 | AAACATCCACCGCGACCTCAAGCC | |
| OsCPK2-F | GGCAACTGCTGTCCTGGCTCT | Detecting nucleotide changes of *OsCPK2* |
| OsCPK2-R | ATGGCGGCAACGGATGAA | |

| | | |
|---|---|---|
| gOsCPK3-F1 | TGTTGGGGAAGTGAGACGACTTGA | Knocking out the endogenous *OsCPK3* gene using the GTTT-type sgRNA scaffold |
| gOsCPK3-R1 | AAACTCAAGTCGTCTCACTTCCCC | |
| gOsCPK3-F2 | GTGTGTCGATGGCCTTGAGCGGCG | Knocking out the endogenous *OsCPK3* gene using the GTTT-type sgRNA scaffold |
| gOsCPK3-R2 | AAACCGCCGCTCAAGGCCATCGAC | |
| gOsCPK3-F1 | TGTTGGGGAAGTGAGACGACTTGA | Knocking out the endogenous *OsCPK3* gene using the GCCC-type sgRNA scaffold |
| gOsCPK3-NG-R1 | GGGCTCAAGTCGTCTCACTTCCCC | |
| gOsCPK3-F2 | GTGTGTCGATGGCCTTGAGCGGCG | Knocking out the endogenous *OsCPK3* gene using the GCCC-type sgRNA scaffold |
| gOsCPK3-NG-R2 | GGGCCGCCGCTCAAGGCCATCGAC | |
| OsCPK3-F1 | CAATTCCTTCTCCTCCCATC | Detecting nucleotide changes of *OsCPK3* |
| OsCPK3-R1 | GTACGTCACCCCGAACTCCC | |
| OsCPK3-F2 | GCAAGTCCATCTCCAAGCGG | |
| OsCPK3-R2 | GTATTCCACCCAAAACAGACCA | |
| gOsCPK4-F1 | TGTTGTCTCATCCCACACTGCGAC | Knocking out the endogenous *OsCPK4* gene using the GTTT-type sgRNA scaffold |
| gOsCPK4-R1 | AAACGTCGCAGTGTGGGATGAGAC | |
| gOsCPK4-F2 | GTGTGCCGTCAAGCGCATCGACAA | Knocking out the endogenous *OsCPK4* gene using the GTTT-type sgRNA scaffold |
| gOsCPK4-R2 | AAACTTGTCGATGCGCTTGACGGC | |
| gOsCPK4-F1 | TGTTGTCTCATCCCACACTGCGAC | Knocking out the endogenous *OsCPK4* gene using |

| | | |
|---|---|---|
| gOsCPK4-NG-R1 | GGGCGTCGCAGTGTGGGATGAGAC | the GCCC-type sgRNA scaffold |
| gOsCPK4-F2 | GTGTGCCGTCAAGCGCATCGACAA | Knocking out the endogenous *OsCPK4* gene using the GCCC-type sgRNA scaffold |
| gOsCPK4-NG-R2 | GGGCTTGTCGATGCGCTTGACGGC | |
| OsCPK4-F | GACCAAACAACCTCCCTC | Detecting nucleotide changes of *OsCPK4* |
| OsCPK4-R | TTCGCACTAGCTCAATCC | |
| gOsCPK5-F3 | TGTTGAGTACTTCGCCAGCTTCCG | Knocking out the endogenous *OsCPK5* gene using the GTTT-type sgRNA scaffold |
| gOsCPK5-R3 | AAACCGGAAGCTGGCGAAGTACTC | |
| gOsCPK5-F4 | GTGTGGAGCGCGTAGTGCTCGCTC | Knocking out the endogenous *OsCPK5* gene using the GTTT-type sgRNA scaffold |
| gOsCPK5-R4 | AAACGAGCGAGCACTACGCGCTCC | |
| OsCPK5-F | TCCAACTCCCTCCATTGCTC | Detecting nucleotide changes of *OsCPK5* |
| OsCPK5-R | ATGTGAACGTACTGCGGGTC | |
| gOsCPK8-F4 | TGTTGCGAACCCCTTCACCGTCGC | Knocking out the endogenous *OsCPK8* gene using the GTTT-type sgRNA scaffold |
| gOsCPK8-R4 | AAACGCGACGGTGAAGGGGTTCGC | |
| gOsCPK8-F5 | GTGTGTCGATATCGACTTGCACGC | Knocking out the endogenous *OsCPK8* gene using the GTTT-type sgRNA scaffold |
| gOsCPK8-R5 | AAACGCGTGCAAGTCGATATCGAC | |
| OsCPK8-F | AACTCACACGGCGAAGCAC | Detecting nucleotide changes of *OsCPK8* |
| OsCPK8-R | ACAGTCCACACGGAGACAC | |

| | | |
|---|---|---|
| gOsCPK14-F1 | TGTTGGATCGACTTGACGTCCTCC | Knocking out the endogenous *OsCPK14* gene using the GTTT-type sgRNA scaffold |
| gOsCPK14-R1 | AAACGGAGGACGTCAAGTCGATCC | |
| gOsCPK14-F2 | GTGTGAGGCGGCGTGCTCGGTGTA | Knocking out the endogenous *OsCPK14* gene using the GTTT-type sgRNA scaffold |
| gOsCPK14-R2 | AAACTACACCGAGCACGCCGCCTC | |
| gOsCPK14-F1 | TGTTGGATCGACTTGACGTCCTCC | Knocking out the endogenous *OsCPK14* gene using the GCCC-type sgRNA scaffold |
| gOsCPK14-NG-R1 | GGGCGGAGGACGTCAAGTCGATCC | |
| gOsCPK14-F2 | GTGTGAGGCGGCGTGCTCGGTGTA | Knocking out the endogenous *OsCPK14* gene using the GCCC-type sgRNA scaffold |
| gOsCPK14-NG-R2 | GGGCTACACCGAGCACGCCGCCTC | |
| OsCPK14-F | AACCGAACCCAAACCCGC | Detecting nucleotide changes of *OsCPK14* |
| OsCPK14-R | GAAGTCGGTGGCCTTGAGAG | |
| gOsCPK20-F1 | TGTTGCGAGTACAACCGGTCGTCG | Knocking out the endogenous *OsCPK20* gene using the GTTT-type sgRNA scaffold |
| gOsCPK20-R1 | AAACCGACGACCGGTTGTACTCGC | |
| gOsCPK20-F2 | GTGTGATAAGCATGGAGTGATGCA | Knocking out the endogenous *OsCPK20* gene using the GTTT-type sgRNA scaffold |
| gOsCPK20-R2 | AAACTGCATCACTCCATGCTTATC | |
| gOsCPK20-F1 | TGTTGCGAGTACAACCGGTCGTCG | Knocking out the endogenous *OsCPK20* gene using the GCCC-type sgRNA scaffold |
| gOsCPK20-NG-R1 | GGGCCGACGACCGGTTGTACTCGC | |
| gOsCPK20-F2 | GTGTGATAAGCATGGAGTGATGCA | Knocking out the endogenous *OsCPK20* gene |

| | | |
|---|---|---|
| gOsCPK20-NG-R2 | GGGCTGCATCACTCCATGCTTATC | using the GCCC-type sgRNA scaffold |
| OsCPK20-F1 | CTCTCGCCTCCTTCTCCT | Detecting nucleotide changes of *OsCPK20* |
| OsCPK20-R1 | GGCATTGTCGTCCTCGTAGGTGT | |
| OsCPK20-F2 | ATCGAGTTCACTGGTGGAGA | |
| OsCPK20-R2 | TGACAACTGAGAACTGACTGAC | |
| gOsCPK21-F1 | TGTTGATCAGCAAGATCTCCCTCG | Knocking out the endogenous *OsCPK21* gene using the GTTT-type sgRNA scaffold |
| gOsCPK21-R1 | AAACCGAGGGAGATCTTGCTGATC | |
| gOsCPK21-F2 | GTGTGTGTGCTTCCGGATCGTCTT | Knocking out the endogenous *OsCPK21* gene using the GTTT-type sgRNA scaffold |
| gOsCPK21-R2 | AAACAAGACGATCCGGAAGCACAC | |
| OsCPK21-F | ACACACACAGAGGAGGAG | Detecting nucleotide changes of *OsCPK21* |
| OsCPK21-R | AATGGGAGAAGAAGAAGA | |
| gOsCPK27-F1 | TGTTGGGAGGCCGTTCTTGGCGAA | Knocking out the endogenous *OsCPK27* gene using the GTTT-type sgRNA scaffold |
| gOsCPK27-R1 | AAACTTCGCCAAGAACGGCCTCCC | |
| gOsCPK27-F2 | GTGTGGCTTGATGTGCGCCGGCTT | Knocking out the endogenous *OsCPK27* gene using the GTTT-type sgRNA scaffold |
| gOsCPK27-R2 | AAACAAGCCGGCGCACATCAAGCC | |
| OsCPK27-F | GAGCGGTGTTGTGTGAAG | Detecting nucleotide changes of *OsCPK27* |
| OsCPK27-R | CGGATGGAGATGATGTTG | |

| | | |
|---|---|---|
| gOsCPK28-F1 | TGTTGCGAGTACGCGTGCAAGTCC | Knocking out the endogenous *OsCPK28* gene using the GTTT-type sgRNA scaffold |
| gOsCPK28-R1 | AAACGGACTTGCACGCGTACTCGC | |
| gOsCPK28-F2 | GTGTGGCGACGATGCGGTCGAAGA | Knocking out the endogenous *OsCPK28* gene using the GTTT-type sgRNA scaffold |
| gOsCPK28-R2 | AAACTCTTCGACCGCATCGTCGCC | |
| gOsCPK28-F1 | TGTTGCGAGTACGCGTGCAAGTCC | Knocking out the endogenous *OsCPK28* gene using the GCCC-type sgRNA scaffold |
| gOsCPK28-NG-R1 | GGGCGGACTTGCACGCGTACTCGC | |
| gOsCPK28-F2 | GTGTGGCGACGATGCGGTCGAAGA | Knocking out the endogenous *OsCPK28* gene using the GCCC-type sgRNA scaffold |
| gOsCPK28-NG-R2 | GGGCTCTTCGACCGCATCGTCGCC | |
| OsCPK28-F | GATCGGAAAATGCAGCCTGA | Detecting nucleotide changes of *OsCPK28* |
| OsCPK28-R | CAAAGTGCGCAGAATCCGAG | |
| gOsMPK3-F1 | TGTTGCAAGCCCATCGGGCGAGGA | Knocking out the endogenous *OsMPK3* gene using the GTTT-type sgRNA scaffold |
| gOsMPK3-R1 | AAACTCCTCGCCCGATGGGCTTGC | |
| gOsMPK3-F2 | TGTTGCTGAAACTTCTCCGGCATC | Knocking out the endogenous *OsMPK3* gene using the GTTT-type sgRNA scaffold |
| gOsMPK3-R2 | AAACGATGCCGGAGAAGTTTCAGC | |
| OsMPK3-F | ACCATAGCCGAGCAACTGAA | Detecting nucleotide changes of *OsMPK3* |
| OsMPK3-R | AAGCAATCATCAAGGGCCA | |
| gOsMPK4-F3 | GTGTGTGACACCAAGTATGTGCCA | Knocking out the endogenous *OsMPK4* gene |

| | | |
|---|---|---|
| gOsMPK4-R3 | AAACTGGCACATACTTGGTGTCAC | using the GTTT-type sgRNA scaffold |
| gOsMPK4-F4 | GTGTGCTCCTCTATAAACCGTGCA | Knocking out the endogenous *OsMPK4* gene using the GTTT-type sgRNA scaffold |
| gOsMPK4-R4 | AAACTGCACGGTTTATAGAGGAGC | |
| OsMPK4-F | TGCATTTTCTCCCCTTCCTGATT | Detecting nucleotide changes of *OsMPK4* |
| OsMPK4-R | ATACGTCATGGTGGAACCGCT | |
| gOsCOI2-F4 | GTGTGAGCGCGTCGACCCTGCACC | Base editing the endogenous *OsCOI2* gene with an NGT PAM |
| gOsCOI2-R4 | AAACGGTGCAGGGTCGACGCGCTC | |
| gOsCOI2-F7 | GTGTGTCGACCCTGCACCAGTGGC | Base editing the endogenous *OsCOI2* gene with an NGC PAM |
| gOsCOI2-R7 | AAACGCCACTGGTGCAGGGTCGAC | |
| gOsCOI2-F8 | GTGTGGAGCGCGTCGACCCTGCAC | Base editing the endogenous *OsCOI2* gene with an NAG PAM |
| gOsCOI2-R8 | AAACGTGCAGGGTCGACGCGCTCC | |
| OsCOI2-F1 | CAACTTCCGCTTTTTCCTTG | Detecting nucleotide changes of *OsCOI2* |
| OsCOI2-R1 | TTGAACGAGGAGAGCATGTG | |
| gOsMPK13-F16 | GTGTGGGACATGGAGTTCTTTACG | Base editing the endogenous *OsMPK13* gene with an NAA PAM |
| gOsMPK13-R16 | AAACCGTAAAGAACTCCATGTCCC | |
| OsMPK13-F1 | TGTGTGCCATTACAGTTTCCA | Detecting nucleotide changes of *OsMPK13* |
| OsMPK13-R1 | CCTGAACTCCCTTCGGGTAG | |

| | | |
|---|---|---|
| gBSRK1-F5 | GTGTGTTCCTAGAACAAACTACAC | Gene correction of the defective endogenous *BSR-K1* gene with an NGA PAM |
| gBSRK1-R5 | AAACGTGTAGTTTGTTCTAGGAAC | |
| gBSRK1-F6 | GTGTGTCCTAGAACAAACTACACT | Gene correction of the defective endogenous *BSR-K1* gene with an NAG PAM |
| gBSRK1-R6 | AAACAGTGTAGTTTGTTCTAGGAC | |
| OsBsr-kit-F1 | GTGAGATGCAAAGCTCGTTGG | Detecting nucleotide changes of *BSR-K1* |
| OsBsr-kit-R1 | CAGGGTGTTGAACAGTTCCG | |
| gMPK8-2 off-1-F2 | AACCTGGTGGAGGTCTGTATC | Detecting the potential off-target mutation of *OsMPK8*-targeting sgRNA 1 |
| gMPK8-2 off-1-R2 | CAGCTGCTTAGCTCTTGGGG | |
| gMPK10-3 off-1-F1 | TCTTAACAGTGGCACCAGGGA | Detecting the potential off-target mutation of *OsMPK10*-targeting sgRNA 2 |
| gMPK10-3 off-1-R1 | CGGCAGCAACAATACCGTTAAG | |
| gMPK10-3 off-2-F1 | TATGCAATCCAGTCATCAAACTCA | Detecting the potential off-target mutation of *OsMPK10*-targeting sgRNA 2 |
| gMPK10-3 off-2-R1 | GCTCATGTGTGGGGTTTGTAG | |
| gMPK10-3 off-3-F1 | AGGTTATGCGGCAGAAATGT | Detecting the potential off-target mutation of *OsMPK10*-targeting sgRNA 2 |
| gMPK10-3 off-3-R1 | TCAGAGGATTTCGAGTGCCG | |
| gCPK3-2 off-1-F1 | GTTATGGCGAGCTAAGGTACG | Detecting the potential off-target mutation of *OsCPK3*-targeting sgRNA 2 |
| gCPK3-2 off-1-R1 | CAACATCGCGGAGTTCAGGG | |
| gCPK3-2 off-2-F2 | ATATACCTCCCCAGAATGGAGGG | Detecting the potential off-target mutation of |

| | | |
|---|---|---|
| gCPK3-2 off-2-R2 | CGAACACTACACATCATCTTCTTC | *OsCPK3*-targeting sgRNA 2 |
| OsCPK1-F | GTCACCTACCTCGTCACCCA | Detecting the potential off-target mutation of *OsCPK3*-targeting sgRNA 2 |
| OsCPK1-R | TTTCAGACCTGTCAAACCCA | |
| gCPK3-2 off-4-F3 | GAGCTGTTCGACCGGATCG | Detecting the potential off-target mutation of *OsCPK3*-targeting sgRNA 2 |
| gCPK3-2 off-4-R3 | GTTCAAGCGTGCATGCATTG | |
| gCPK3-2 off-5-F2 | AAGCGGAAGCTACTCACCGA | Detecting the potential off-target mutation of *OsCPK3*-targeting sgRNA 2 |
| gCPK3-2 off-5-R2 | GCCTACCTACCTGCCCAGAA | |
| gCPK3-2 off-6-F1 | CACCTCTCCGGCCAGCCA | Detecting the potential off-target mutation of *OsCPK3*-targeting sgRNA 2 |
| gCPK3-2 off-6-R1 | GTCCGCGAAGGATGGCGGT | |
| gCPK28-2 off-1-F1 | CATGGAGAGGCCGAAATCAGT | Detecting the potential off-target mutation of *OsCPK28*-targeting sgRNA 1 |
| gCPK28-2 off-1-R1 | AGAAGCTTGGGTGTCCGTAG | |
| gCPK28-1 off-2-F1 | CGAACACCACCCAGCAGCCT | Detecting the potential off-target mutation of *OsCPK28*-targeting sgRNA 1 |
| gCPK28-1 off-2-R1 | CGCTCCGTGTAGTGCCCCTT | |
| gCPK28-2 off-1-F1 | CATGGAGAGGCCGAAATCAGT | Detecting the potential off-target mutation of *OsCPK28*-targeting sgRNA 2 |
| gCPK28-2 off-1-R1 | AGAAGCTTGGGTGTCCGTAG | |
| OsCPK29-F1 | GCGTGCAAGTCGATCAGCAA | Detecting the potential off-target mutation of *OsCPK28*-targeting sgRNA 2 |
| OsCPK29-R1 | GCATGAGCATGTGGGTGGTG | |

| | | |
|---|---|---|
| gCPK28-2 off-4-F1 | GCCAAAATCGGTGGCTTTGA | Detecting the potential off-target mutation of *OsCPK28*-targeting sgRNA 2 |
| gCPK28-2 off-4-R1a | GCTACGCCTGCAAGTCCATC | |
| gCPK28-2 off-5-F3 | GTGAGTTCCTTCCTACCTGG | Detecting the potential off-target mutation of *OsCPK28*-targeting sgRNA 2 |
| gCPK28-2 off-5-R3 | GAGGTGGCAGTACAGGAGAT | |
| gCPK28-2 off-6-F1 | TTCAAAGAGGCAGGCATGGA | Detecting the potential off-target mutation of *OsCPK28*-targeting sgRNA 2 |
| gCPK28-2 off-6-R1 | CTCCAAGAAGAAGCTCCGCA | |
| OsCPK3-F2 | GCAAGTCCATCTCCAAGCGG | Detecting the potential off-target mutation of *OsCPK28*-targeting sgRNA 2 |
| OsCPK3-R2 | GTATTCCACCCAAAACAGACCA | |
| OsCPK1-F | GTCACCTACCTCGTCACCCA | Detecting the potential off-target mutation of *OsCPK28*-targeting sgRNA 2 |
| OsCPK1-R | TTTCAGACCTGTCAAACCCA | |

**Table S3. Rice genes for targeted genome editing in this study.**

| Gene Name | Gene Identifier | Plasmid | Application |
|---|---|---|---|
| *OsMPK3* | Os02g0148100 | pUbi:SpRY | Gene knockout |
| *OsMPK4* | Os06g0699400 | pUbi:SpRY | Gene knockout |
| *OsMPK8* | Os01g0665200 | pUbi:SpRY | Gene knockout |
| *OsMPK9* | Os05g0582400 | pUbi:SpRY | Gene knockout |
| *OsMPK10* | Os01g0629900 | pUbi:SpRY | Gene knockout |
| *OsMPK13* | Os02g0135200 | pUbi:rBE62 | Adenine base editing |
| *OsCPK1* | Os01g0622600 | pUbi:SpRY | Gene knockout |
| *OsCPK2* | Os01g0808400 | pUbi:SpRY | Gene knockout |
| *OsCPK3* | Os01g0832300 | pUbi:SpRY | Gene knockout |
| *OsCPK4* | Os02g0126400 | pUbi:SpRY | Gene knockout |
| *OsCPK5* | Os02g0685900 | pUbi:SpRY | Gene knockout |
| *OsCPK8* | Os03g0808600 | pUbi:SpRY | Gene knockout |
| *OsCPK14* | Os05g0491900 | pUbi:SpRY | Gene knockout |
| *OsCPK20* | Os07g0568600 | pUbi:SpRY | Gene knockout |
| *OsCPK21* | Os08g0540400 | pUbi:SpRY | Gene knockout |
| *OsCPK27* | Os12g0486600 | pUbi:SpRY | Gene knockout |
| *OsCPK28* | Os12g0169800 | pUbi:SpRY | Gene knockout |
| *OsCERK1* | Os08g0538300 | pUbi:SpG | Gene knockout |
| *OsGSK4* | Os06g0547900 | pUbi:SpG | Gene knockout |
| *OsETR2* | Os04g0169100 | pUbi:SpG | Gene knockout |
| *OsCOI2* | Os03g0265500 | pUbi:rBE66 | Cytosine base editing |
| *OsGS1* | Os02g0735200 | pUbi:rBE62 | Adenine base editing |
| *BSR-K1* | Os10g0548200 | pUbi:rBE66 | Cytosine base editing |
